# scDisentangle: sequential disentanglement for single-cell perturbation response prediction

**DOI:** 10.64898/2026.11.04.764292

**Authors:** Salim El Mejjad, Salwa Enezari, Anas Bedraoui, Fatima Zahra El Hajji, Rachid El Fatimy, Matan Hofree, Tariq Daouda

## Abstract

Predicting the transcriptional response of single cells to perturbation is a central goal of single-cell modeling, with broad application to analysis and experimental design. Existing methods often fail to beat simple baselines and offer little insight into how biological variation is organized. We present scDisentangle, a model that decomposes single-cell expression into observed attributes and a sequence of residual components, generating counterfactuals that preserve source cell identity. scDisentangle outperformed current models and naive baselines on cytokine stimulation, liver infection, and unseen combinatorial genetic perturbations, on which accuracy was largely independent of how closely a response resembled the average perturbation effect. In an immune checkpoint inhibitor-associated myocarditis cohort, cell-resolved counterfactuals predicted donor-specific intracardiac CD8+ T cell states from peripheral blood. In a castration-regeneration prostate atlas, predictions recapitulated luminal dynamics and progenitor-marker kinetics upon androgen add-back. scDisentangle’s coarse-to-fine latent structure aligned with hematopoietic lineage organization, with early levels capturing a less differentiated core progressively refined into cell-type-specific programs.

## Introduction

Predicting cellular responses to perturbations is central to understanding and engineering cell state. Single-cell RNA sequencing (scRNA-seq) provides the resolution to capture these responses, yet cells and their perturbation responses are inherently heterogeneous, and the destructive nature of scRNA-seq precludes observing the same cell in multiple states. Computational models are therefore needed to infer perturbation responses from unpaired measurements. scGen^1^ and scVIDR^2^ predict perturbation responses through vector arithmetic in latent space, but assume homogeneous perturbation effects across all cells or within predefined cell types.

Disentangled representation learning methods instead separate perturbation effects from other sources of variation, allowing for cell-specific counterfactual predictions. Nonetheless, adversarial^3,4^ and distributional-matching^5^ approaches risk removing context-dependent biological variation, while encoder-less approaches^6^ remain susceptible to entanglement through the reconstruction objective.

Recent systematic evaluations showed that current models fail to consistently outperform simple baselines^7,8^, raising doubts about what perturbation-specific signal they capture beyond shared control-to-perturbed shifts. A further limitation lies in evaluation: current metrics operate on population-level averages and overlook whether predictions preserve cell identity. To directly address this concern, we propose an identity preservation metric that tests whether a predicted perturbation profile for a control cell remains closer to its source cell than to other controls.

Here, we present scDisentangle (Fig. 1), a generative model that decomposes each cell into attribute-related embeddings and a sequence of residual factors. Each successive factor captures the residual variation left unexplained by attributes and earlier factors, and as a result information shared between the attribute embeddings and the residual factors is suppressed by construction. This is done without computationally demanding adversarial training or distributional matching. Additionally, this process lends itself naturally to generation of counterfactual predictions, which are obtained by substituting attribute-related embeddings at inference time while preserving cell-specific residual factors.

**Fig. 1:**
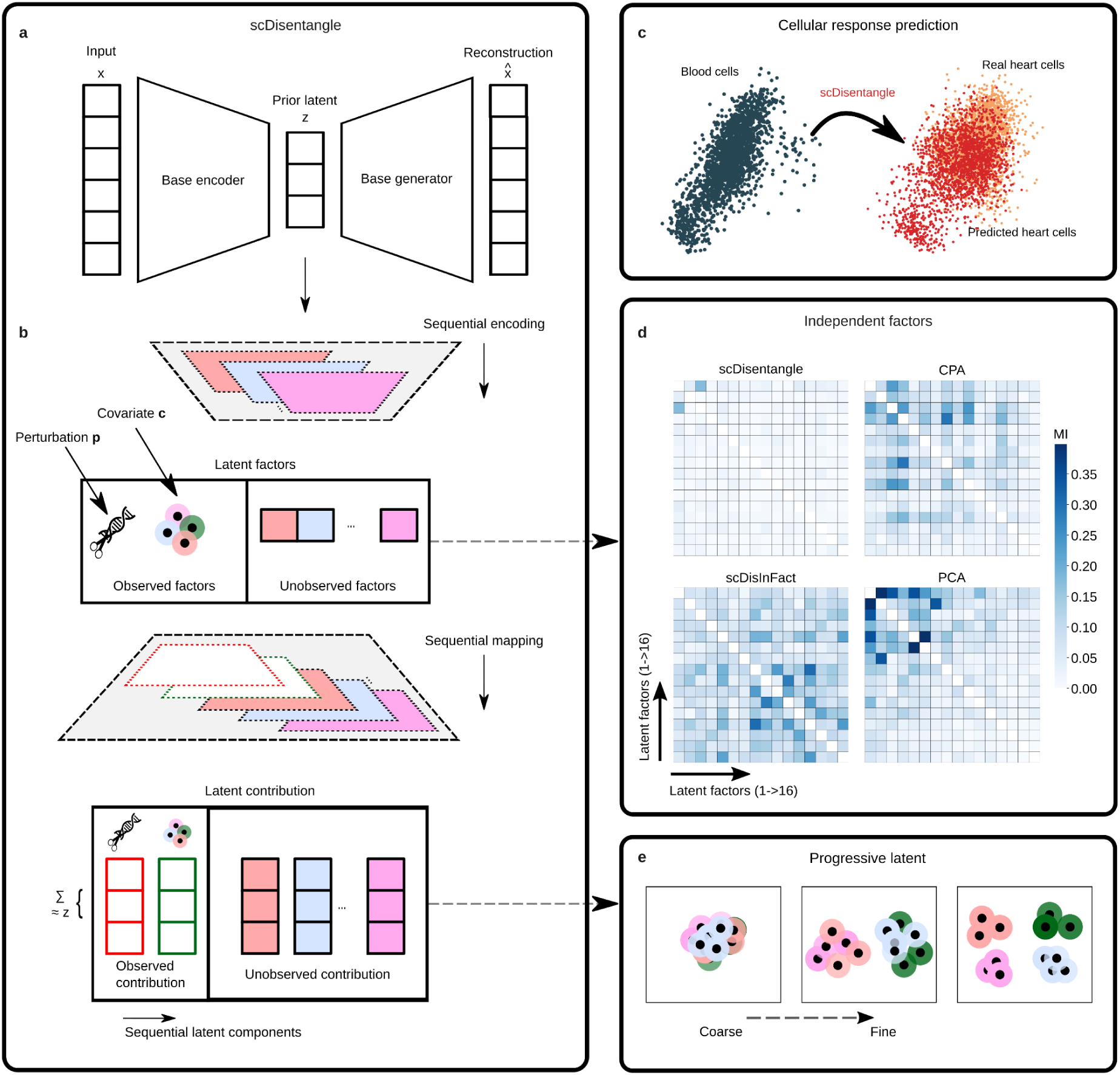
scDisentangle framework for capturing complementary variation by sequential decomposition of known and latent factors. **a,** Base autoencoder. The base encoder maps log-transformed gene expression to an initial latent representation z; the base generator decodes z back to gene expression space. **b,** Sequential decomposition. Each observed covariate is embedded (with sine activation) and projected to the dimensionality of z alongside an encoded class-mean baseline. A series of encoder-mapper pairs iteratively extracts residual variation in z into 16 scalar latent factors; at each step, the accumulated factors are mapped to the dimensionality of z. All contributions are summed to recover the initial latent representation. **c,** Illustration of counterfactual generation. Given control (blood) CD8+ T cells and a target perturbation (heart tissue), scDisentangle generates the corresponding heart-infiltrating CD8+ T expression profiles (Point clouds adapted from Fig. 3a). **d,** Pairwise mutual information between latent factors learned by scDisentangle, CPA, scDisInFact, and between principal components of PCA on a PBMC dataset. Latent dimensionality was matched across methods (16 dimensions; Extended Data Fig. 8). **e,** Illustration of the progressive latent structure: Earlier levels capture broad variation while later levels capture finer biological distinctions.

Across benchmarks involving immune stimulation, pathogen infection, and combinatorial genetic perturbation, scDisentangle outperforms competing methods and the naive baselines recently shown to rival them^8^. In paired blood and cardiac tissue, scDisentangle predicts transcriptional shifts of heart-infiltrating CD8+ T cells beyond population-level baselines. Moreover, unlike competing methods, scDisentangle shows weaker dependence on shared perturbation signatures. In a mouse prostate atlas spanning a castration-regeneration cycle, it reproduces luminal and progenitor marker dynamics and predicts cell-state transitions across the cycle. scDisentangle’s progressive factors also recapitulate hematopoietic lineage organization across two PBMC datasets and T cell subset structure in B-cell acute lymphoblastic leukemia.

## Results

### Model overview

scDisentangle takes single-cell expression data with annotated attributes such as stimulation status and cell type, and encodes each cell into an initial latent representation (Fig. 1a), which is decomposed into attribute-related contributions and a sequence of latent factor contributions, each produced by its own encoder (Fig. 1b). Counterfactual predictions are generated by replacing the embedding and encoded class-mean baseline for attributes of interest while keeping cell-specific factors fixed and decoding the recomposed representation into the predicted single-cell expression profile (Fig. 1c). Successive factors capture complementary variation rather than redundant information, without adversarial training or explicit independence penalties (Fig. 1d). Finally, the training yields a series of coarse-to-fine representations that can be examined level by level, with earlier levels capturing broader axes of variation and subsequent levels capturing finer levels of cellular heterogeneity (Fig. 1e), thus aiding interpretation.

### scDisentangle predicts cell type-specific IFN-β responses

We benchmarked scDisentangle on a PBMC dataset^9^ (Supplementary Table 1), comprising eight immune cell types profiled under IFN-β stimulation and control. We defined out-of-distribution (OOD) settings as excluding stimulated cells from one cell type during training, then predicting their profiles from control cells (10 seeds per setting). We compared scDisentangle to four competing methods: scGen^1^, CPA^3^, biolord^6^, and scDisInFact^5^. To address recent concerns about deep learning models not consistently outperforming simple baselines^7,8^, we also evaluated two naive baselines: a perturbation-agnostic baseline (mean control expression of the held-out cell type) and a context-agnostic baseline (mean stimulated expression across training cells). scDisentangle consistently and significantly outperformed all competitors and both baselines where applicable across correlation (Fig. 2a; Extended Data Fig. 1a,b), distributional (Fig. 2b), and error metrics (Extended Data Fig. 1c,d; Supplementary Table 2), evaluated on top differentially expressed genes (DEGs). scDisentangle also achieved the highest source-cell proximity score, suggestive of improved identity preservation (Fig. 2c) and its predictions for NK cells matched the observed distribution of the top IFN-β-upregulated gene (ISG15; Fig. 2d; Supplementary Fig. 1), illustrating that the predicted population can reproduce cell-to-cell expression variation for a representative response gene.

**Fig. 2:**
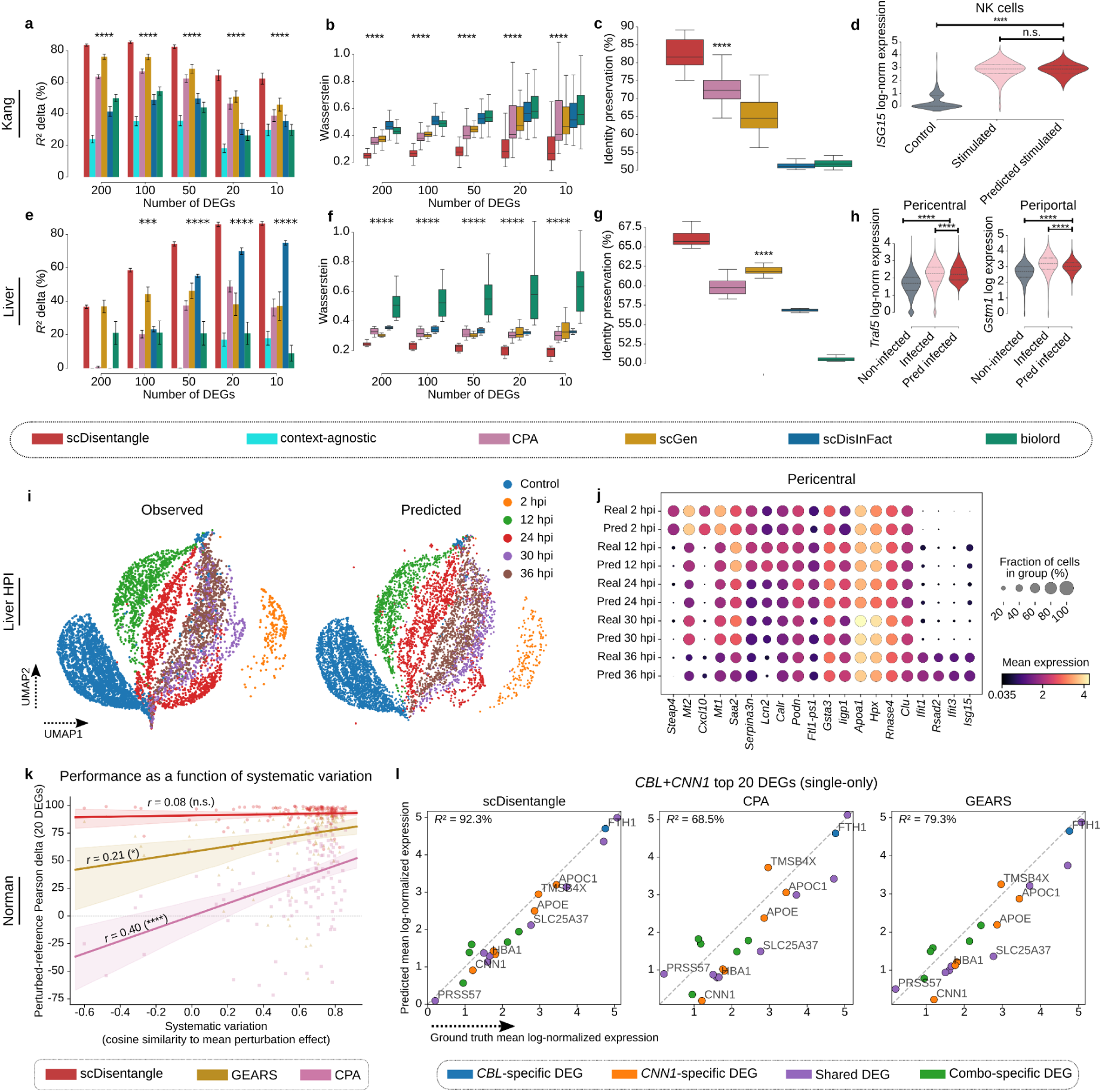
scDisentangle outperforms state-of-the-art perturbation models. **a-c,** Benchmark of scDisentangle against CPA, scGen, scDisInFact, biolord (**a**-**c**), and the context-agnostic baseline (which corresponds to the mean stimulated cells in the training-set; **a**) on *R*² delta (**a**), Wasserstein (**b**) and identity preservation (**c**); *n* = 80 (8 cell types x 10 seeds), Kang. **d,** Violin plot of real control, real stimulated, and scDisentangle-predicted stimulated cells for the top differentially expressed gene (DEG; stimulated vs. control) in NK cells (seed = 1). Horizontal lines: median and quartiles. Significance: two-sided t-tests, unpaired (real stimulated vs. predicted stimulated) and paired (control vs. predicted stimulated). **e-g,** Benchmark on Liver using *R*² delta (**e**), Wasserstein (**f**) and identity preservation (**g**); *n* = 20 (2 zones x 10 seeds). **h,** Violin plots of non-infected, infected and scDisentangle-predicted infected cells for the top DEG (infected vs. non-infected) in pericentral (*Traf5*; left) and periportal (*Gstm1*; right) zones (seed = 1). Lines and statistical significance as in **d**. **i,** UMAP of observed and scDisentangle-predicted expression states for held-out infected cells, embedded jointly with observed data across both spatial zones, coloured by infection time point. **j,** Dot plot comparing observed vs. scDisentangle-predicted expression of infection time-specific marker genes (pericentral zone) across the time course. **k,** Performance vs. systematic variation across 128 Norman combinatorial perturbations, y-axis: perturbed-reference Pearson delta on top 20 DEGs. Lines: linear fits, *r*: Pearson correlation between systematic variation and performance (two-sided Pearson test). Bands: bootstrapped 95% CI (2,000 resamples). **l,** Predicted vs. observed mean expression for the *CBL*+*CNN1* combinatorial perturbation (single-only out-of-distribution scenario). Scatter plots across the top 20 DEGs for scDisentangle (left), CPA (middle), and GEARS (right). Genes coloured by DEG membership. Dashed line: identity (y = x). *R*² computed between the 20-gene ground-truth and predicted means. Neither *CBL* nor *CNN1* was seen in combination during training. Bars (**a**,**e**): mean ± s.e.m. Boxplots (**b**,**c**,**f**,**g**), centre: median; box: first and third quartiles; whiskers: 1.5× IQR; outliers not shown. For **a**-**c** and **e**-**g**, significance was assessed using one-sided Mann-Whitney U tests against the best-performing competing method and context-level p-values were combined using Fisher’s method. For **d**, **h**, and **k**, n.s. indicates not significant. \**P* < 0.05, \*\**P* < 0.01, \*\*\**P* < 0.001, \*\*\*\**P* < 0.0001. Expression values are log-normalized.

### scDisentangle predicts spatio-temporal liver infection responses

We next benchmarked scDisentangle on a mouse liver dataset^10^ (Supplementary Table 1) of hepatocytes from Plasmodium-injected and control mice, annotated by spatial zonation (periportal or pericentral). In each OOD setting, we excluded infected cells from one zone during training and predicted their transcriptional profiles from non-infected cells of that zone (10 seeds per setting). scDisentangle surpassed competing methods under their recommended settings^1,3,5,6^, and naive baselines where applicable, across evaluated metrics (Fig. 2e,f; Extended Data Fig. 2; Supplementary Table 3), and achieved the highest identity preservation (Fig. 2g). scDisentangle also recapitulated the distributional shift of top infection-responsive DEGs in both pericentral (*Traf5*) and periportal (*Gstm1*) cells (Fig. 2h).

We then asked whether scDisentangle captures spatio-temporal infection dynamics. We withheld infected cells from one zone across all five post-infection time points and inferred their expression profiles by applying infection status and time-point covariates to control cells of the same zone. scDisentangle’s predictions largely aligned with observed infected cells across the infection time course (Fig. 2i), recapitulating the levels and fractions of expressing cells for time-point-specific DEGs (Fig. 2j; Extended Data Fig. 3). scDisentangle therefore reproduced selected temporal expression patterns across the infection time course.

### scDisentangle predicts responses to unseen combinations of genetic perturbations

Genes can interact non-additively^11,12^, potentially driving cells into unexpected transcriptional states that cannot be predicted from individual gene effects alone. CRISPR perturbation screens can map these interactions, yet testing every gene pair quickly becomes impractical: 105 genes require 5,460 experiments. We benchmarked scDisentangle on the Perturb-seq CRISPRa dataset of Norman et al^11^, comprising 105 single and 131 combinatorial perturbations (filtered to 101 and 128; Supplementary Tables 1 and 4), and evaluated generalization under two OOD scenarios of increasing difficulty. In the combinatorially-seen scenario, a double perturbation A+B is withheld, but both constituent genes may appear in other combinations during training. In the harder single-only scenario, A and B are observed only as single perturbations, never in any combination. We benchmarked scDisentangle against CPA^3^ and GEARS^13^, and two baselines from Systema^8^: match-mean, which predicts A+B as the average of mean A and mean B across training cells, and perturb-mean, which predicts the response as the mean across perturbed training cells.

In the combinatorially-seen scenario, we followed the CPA benchmarking protocol and trained each model across 13 splits, covering all 128 double perturbations (Supplementary Table 5). scDisentangle outperformed both CPA and GEARS across correlation (Extended Data Fig. 4a-d) and error (Extended Data Fig. 4e,f; Supplementary Table 6) metrics. On *R*² delta, scDisentangle significantly surpassed all methods including match-mean (81.9% vs 67.4% on top-20 DEGs). On Pearson delta, it surpassed both deep learning models and was competitive with match-mean. Consistent with previous reports^8^, match-mean outperformed CPA and GEARS on *R*² delta, but did not match scDisentangle. scDisentangle also achieved the highest specificity scores, including centroid accuracy (97.4% vs 84.6% for match-mean), indicating that predicted profiles reliably mapped to their correct perturbation identity, and achieved the highest identity preservation (Extended Data Fig. 4g-i). Results were consistent when evaluating on combination-specific DEGs: genes differentially expressed in the double perturbation relative to its single constituents (Supplementary Fig. 2a-f; Supplementary Table 7).

A recent systematic evaluation showed that perturbation models struggle to generalize beyond systematic variation, that is, the shared transcriptional shift common to all perturbations^8^. Following this framework, we quantified the systematic variation of each held-out combination as the cosine similarity between its differential expression and the mean perturbation effect. We then tested whether model performance, measured by perturbed-reference Pearson delta (prediction-observation correlation after subtracting the mean perturbed profile) scaled with systematic variation. CPA and GEARS showed positive correlations between performance and systematic variation (*r* = 0.40 and *r* = 0.21, respectively), indicating that both models perform better when a perturbation response resembles the average perturbation effect. By contrast, scDisentangle showed no significant association (*r* = 0.08, *P* = 0.38; Fig. 2k). A similar trend was observed for Pearson delta, with scDisentangle remaining the least correlated with systematic variation (Supplementary Fig. 3).

In the single-only scenario, we selected nine OOD perturbations spanning six non-additive genetic interaction categories^11^ (Supplementary Table 8). For the pair CBL+CNN1, scDisentangle’s predicted expression closely matched observed levels across the top 20 DEGs, including genes unique to the combination, while CPA and GEARS showed larger deviations (Fig. 2l). Similar advantages were observed across other held-out pairs (Extended Data Fig. 5; Supplementary Figs 4 and 5). scDisentangle achieved the highest mean *R*² delta (77.0% vs 67.1% for match-mean, 44.0% for GEARS, and 20.0% for CPA on top-20 DEGs; Extended Data Fig. 6a-d) and the lowest error (Extended Data Fig. 6e,f; Supplementary Table 9). scDisentangle also achieved the highest specificity scores (98.0% vs 80.1% for match-mean on centroid accuracy; Extended Data Fig. 6g,h) and the highest identity preservation (76.0% vs 53.9% for CPA; Extended Data Fig. 6i). The advantage over match-mean was more notable on combination-specific DEGs where scDisentangle achieved 75.3% *R*² delta versus 55.7% for match-mean (Supplementary Fig. 6; Supplementary Table 10; additional metrics in Supplementary Figs 7 and 8). This is expected: match-mean averages single perturbation profiles, and cannot, by construction, capture combination-specific interaction effects.

These results show that scDisentangle captures perturbation-specific transcriptional effects beyond shared perturbation signatures that averaging baselines cannot recover.

### scDisentangle predicts patient-specific transcriptional adaptations of CD8+ T cells upon heart infiltration

As a proof-of-concept for cross-tissue prediction, we applied scDisentangle to model heart-associated transcriptional states of CD8+ T cells in immune checkpoint inhibitor (ICI)-associated myocarditis, a potentially fatal adverse event driven by cytotoxic T cells through incompletely understood patient-specific adaptations. We used a paired peripheral blood and cardiac scRNA-seq dataset^14^ (Supplementary Table 1) from 12 ICI-treated patient donors. We trained scDisentangle in 11 leave-one-donor-out configurations (one donor excluded due to insufficient CD8+ T cells), holding out intracardiac cells from one donor and repeating each configuration across 10 random seeds. Blood cells from each held-out donor were then used to predict their transcriptional state upon heart infiltration (Fig. 3a). We compared predictions for CD8+ T cells with two baselines: a context-agnostic baseline (Heart-mean), and a perturbation-agnostic baseline (Blood-mean). Heart-mean assumes a donor-invariant tissue effect, while Blood-mean assumes no transcriptional change between T cells from blood and heart. scDisentangle significantly outperformed baselines on correlation (Fig. 3b; Supplementary Fig. 9a,b) and error (Fig. 3c; Supplementary Fig. 9c; Supplementary Table 11) metrics, capturing donor-associated differences in heart CD8+ T-cell expression that were not recovered by population-mean blood or heart baselines. Results were consistent when evaluating all cells (Supplementary Fig. 10; Supplementary Table 12). We further compared the predicted expression distributions with those of real circulating and intracardiac CD8+ T cells across all held-out donors (Fig. 3d). scDisentangle reproduced the upregulation of cytotoxicity effector *PRF1*, chemokine *CCL4,* and *MTRNR2L8*, as well as the downregulation of *HLA-B*, *IFITM1*, and *S100A4*.

**Fig. 3:**
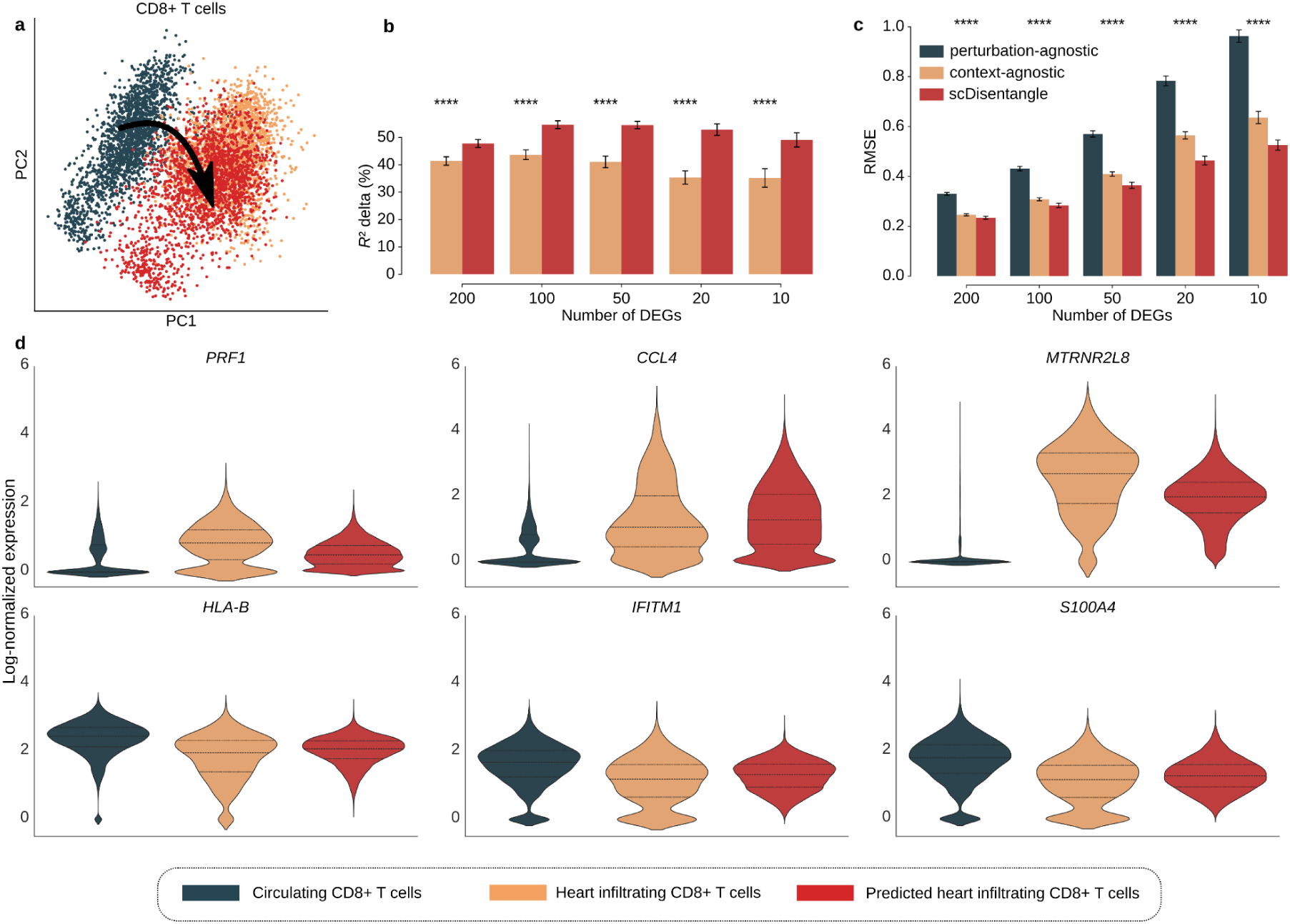
scDisentangle learns patient-specific responses of CD8+ T cells upon heart infiltration. **a,** PCA of real circulating CD8+ T cells (dark teal), real heart-infiltrating CD8+ T cells (orange), and scDisentangle-predicted heart-infiltrating CD8+ T cells (red), generated by counterfactually mapping each donor’s blood cells to heart tissue. Predictions were collected across all 11 OOD donors (seed = 1); data were equally subsampled to 1,975 cells per group for visualization. Arrow illustrates predicted transition from blood to heart. **b,c,** Benchmark of scDisentangle against naive baselines on *R*² delta (**b**) and RMSE (**c**) on Myocarditis (CD8+ T cells subset). context-agnostic corresponds to the CD8+ T mean in heart, while perturbation-agnostic corresponds to the OOD donor’s CD8+ T mean in blood. Bars indicate the mean and error bars the standard error of the mean (s.e.m.) across *n* = 110 independent evaluations (11 OOD donors × 10 random seeds). Significance was assessed using one-sided Mann-Whitney U tests applied within each OOD context against the best-performing baseline at each DEG count; context-level p-values were combined using Fisher’s method; \**P* < 0.05, \*\**P* < 0.01, \*\*\**P* < 0.001, \*\*\*\**P* < 0.0001. **d,** Violin plots comparing single-cell log-normalized expression distributions of *PRF1*, *CCL4*, *MTRNR2L8*, *HLA-B*, *IFITM1*, and *S100A4* (selected from top DEGs) in CD8+ T cells across real circulating, real heart-infiltrating, and scDisentangle-predicted heart-infiltrating CD8+ T cells (seed = 1). Horizontal lines indicate the median and quartiles.

### scDisentangle tracks luminal prostate transcriptional dynamics over the castration-regeneration (C/R) cycle

Androgen deprivation therapy is a first-line treatment for high-risk and advanced prostate cancer^15,16^ and induces prostate gland shrinkage and tumor regression^17^. However, this response is reversible: upon androgen restoration, prostate epithelial cells rapidly re-enter a proliferative state to regenerate the gland^18^. We applied scDisentangle to a mouse prostate scRNA-seq atlas comprising 87,187 cells (Supplementary Table 1) sampled over a full C/R cycle^19^. UMAP^20^ and PCA visualizations of the disentangled representation showed that C/R stage-associated variation was separated from major prostate cell population structure (Fig. 4a,b). We used scDisentangle to model luminal cell dynamics across the C/R cycle. For each C/R stage, we generated counterfactual L2 profiles by applying the corresponding temporal intervention to intact cells. L2 cells from the target stage were excluded during training. We quantified L1-L2 proximity by computing the mean absolute error between L1 and L2 centroids for each C/R stage in a shared PCA space fit on observed cells: scDisentangle captured the L1-L2 convergence post-castration, and divergence following androgen add-back (Fig. 4c). Stage-wise PCA visualizations further recapitulated the co-clustering of L1 and L2 cells at day 28 post-castration and day 1 post-regeneration, followed by their separation at later regeneration stages (Fig. 4d; Supplementary Fig. 11). Karthaus et al.^19^ attributed this convergence to the loss of androgen receptor (AR)-dependent markers such as *Psca* in L2 cells. scDisentangle likewise captured the *Psca* decline during castration and its return toward pre-castration levels after androgen add-back (Fig. 4e). We additionally compared observed and generated expression distributions of three markers: *Pbsn*, *Tgm4* and *Ly6a*. *Pbsn* and *Tgm4* are AR-regulated and typically decline after castration^19,21^, whereas *Ly6a,* whose transcription is not androgen-dependent^19^, remains relatively stable; scDisentangle recapitulated these dynamics (Fig. 4f; Supplementary Fig. 12; additional markers in Supplementary Fig. 13).

**Fig. 4:**
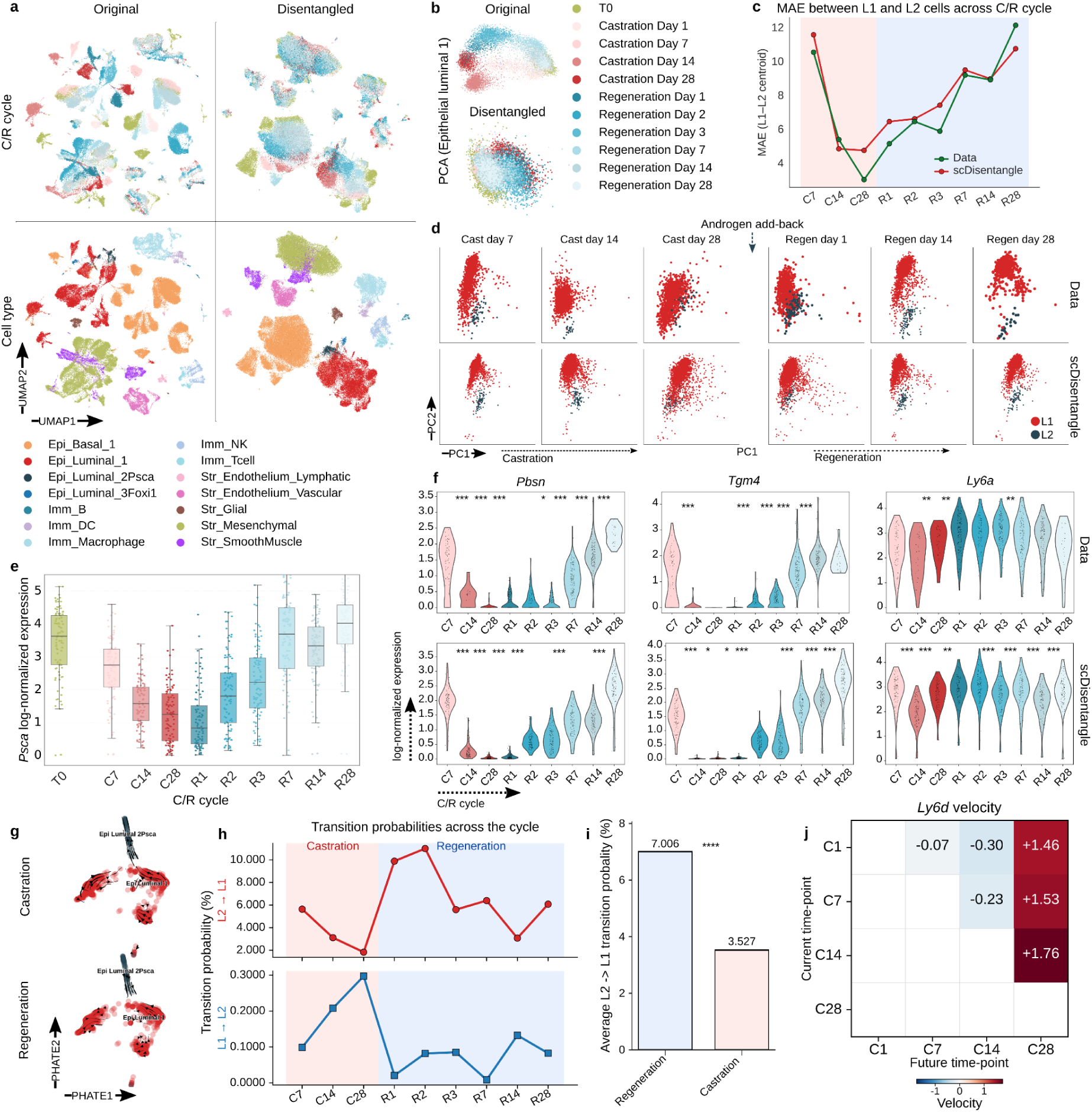
scDisentangle tracks prostate transcriptional dynamics over the castration-regeneration (C/R) cycle. **a,** UMAP of original data, and scDisentangle’s disentangled latent representation, coloured by C/R stage (top) and cell type (bottom). **b,** PCA of disentangled representation on L1 cells coloured by C/R stage. **c,** MAE (mean absolute error) between L1 and L2 centroids in a shared PCA space for observed (green) and scDisentangle-generated (red) cells across the C/R cycle. Stage C1 (*n* = 1 cell) excluded. **d,** PCAs of observed (top) vs. scDisentangle-generated (bottom) L1 and L2 cells. **e,** Boxplot of scDisentangle-generated *Psca* log-normalized expression across C/R stages. Centre line: median; box: first and third quartiles; whiskers: 1.5× IQR; outliers not shown; individual data points overlaid. **f,** Violin plots showing log-normalized expression of *Pbsn*, *Tgm4*, and *Ly6a* in L2 cells across the C/R cycle. Top row: observed expression. Bottom row: scDisentangle-generated expression from control L2 cells counterfactually mapped to each C/R stage. Individual data points overlaid. Significance: t-test, two-sided independent (observed) or paired (scDisentangle-generated) between consecutive stages; \**P* < 0.05, \*\**P* < 0.01, \*\*\**P* < 0.001. **g,** Velocity stream plots of scDisentangle counterfactuals on PHATE embeddings. T0 L1 (red) and L2 (dark teal) cells embedded by PHATE computed on original log-normalized profiles (shared across both). Arrows: inferred direction of transcriptional change. Top: late castration (C7 → C28). Bottom: onset of regeneration (C28 → R1). **h,** Mean per-cell transition probability (%), L2 to L1 (top) and L1 to L2 cells (bottom) inferred by scVelo for each pair of consecutive generated C/R stages. **i,** Mean L2-to-L1 transition probability averaged over castration and regeneration stages. Significance: one-sided Wilcoxon signed-rank test (regeneration > castration); \**P* < 0.05, \*\**P* < 0.01, \*\*\**P* < 0.001, \*\*\*\**P* < 0.0001. **j,** Mean *Ly6d* velocity (future - current counterfactual expression) in L2 cells. Rows: current state; columns: future state.

To characterize luminal lineage dynamics, we repurposed scVelo^22^ to infer cell-state transition probabilities from generated expression profiles. For each pair of consecutive C/R stages, we generated both the current and the next time point from the same set of control L1 and L2 cells, defined the per-cell velocity as future minus current expression, and used scVelo to construct a cell-to-cell transition probability matrix. Velocity stream plots on PHATE^23^ embeddings revealed a net flow from L1 toward L2 during late castration and its reversal at the onset of regeneration (Fig. 4g). Quantitatively, L2-to-L1 transition probability declined progressively during castration and spiked 5.4-fold at the onset of regeneration relative to the preceding transition (Fig. 4h,i; Extended Data Fig. 7a,b), coinciding with the rapid recruitment of luminal cells into the cell cycle upon androgen restoration^19^. Conversely, L1-to-L2 transition probability increased slightly during castration and declined upon androgen restoration, consistent with the convergence of L1 and L2 cells under androgen deprivation (Fig. 4h; Extended Data Fig. 7c,d).

Recently, *Ly6d* was linked to castration-resistant prostate progenitors with multipotent differentiation^24^. *Ly6d*+ cells are enriched after castration and, upon androgen restoration, give rise to *Ly6d*-progeny. scDisentangle-derived velocities revealed a sharp upregulation of *Ly6d* at late castration (day 28; Fig. 4j), followed by a rapid decline upon androgen add-back (Extended Data Fig. 7e,f), recapitulating the dynamics of this progenitor marker.

### scDisentangle learns latent representations that recapitulate hematopoietic lineage organization

Whether latent factors learned by generative models capture interpretable biological variation remains an open question in single-cell analysis. scDisentangle learns complementary variation by design. Consequently, pairwise Pearson correlations between learned factors are lower than those of CPA^3^ and scDisInFact^5^ (Extended Data Fig. 8a,b). While PCA enforces zero linear correlation by construction, scDisentangle shows lower nonlinear dependence, as measured by pairwise mutual information (Extended Data Fig. 8c,d). To interpret how structure is captured across learned factors, we progressively added latent factors, revealing increasingly fine-grained biological structure. On two PBMC datasets from Kang^9^ and Hao^25^, early latent levels capture broad hematopoietic lineage structure, separating myeloid from lymphoid compartments. Subsequent levels progressively refine this separation, distinguishing T cells from other lymphoid populations, resolving myeloid subpopulations, and finally separating T-cell subtypes at the finest levels (Fig. 5a). These patterns recapitulate the coarse-to-fine organization of hematopoietic lineages, while remaining largely invariant to the observed condition (Fig. 5b; Supplementary Fig. 14 and 15). Notably, no progenitor populations were present in either training set.

**Fig. 5:**
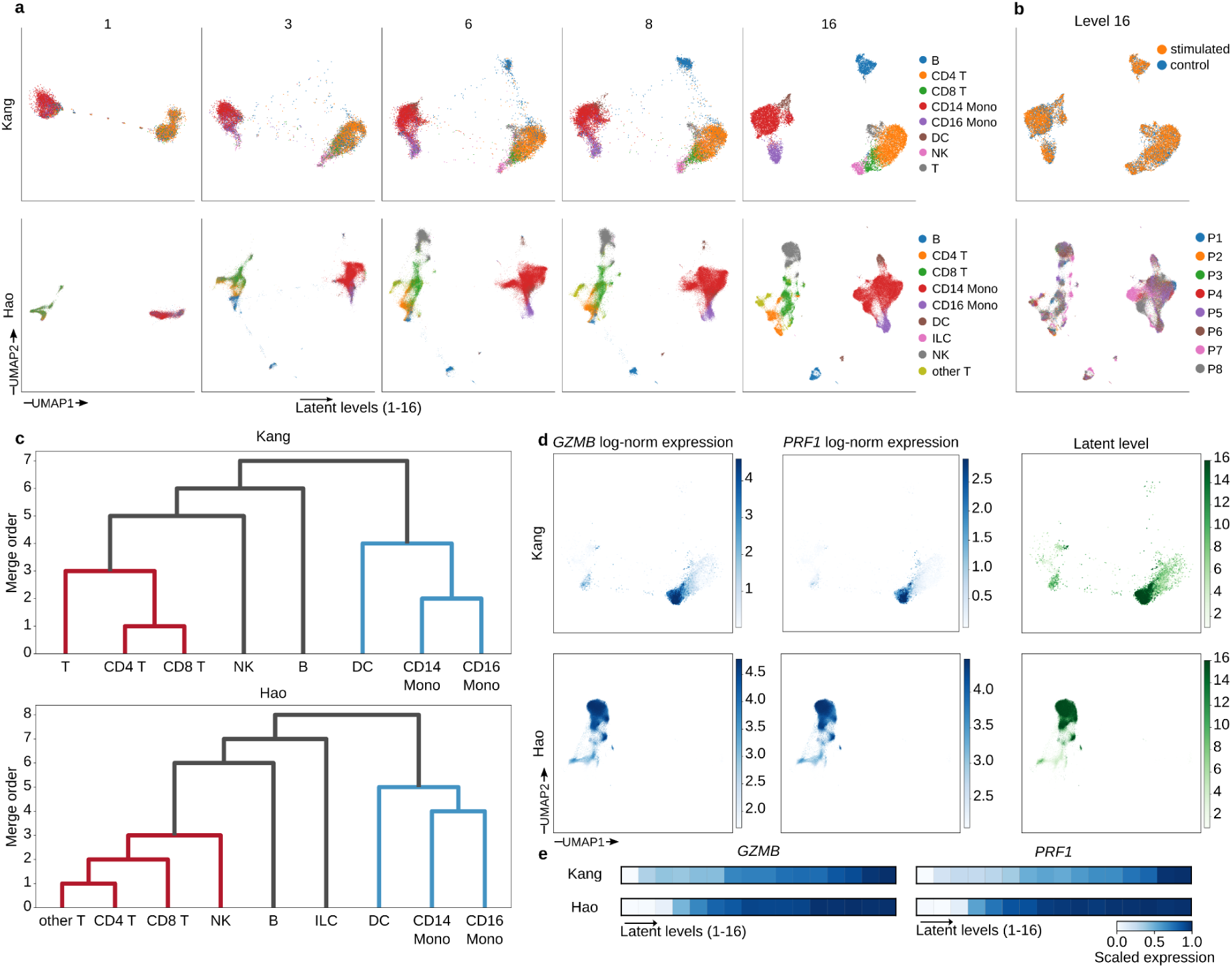
scDisentangle learns structured latent representations that recapitulate hematopoietic lineage organization. **a,** UMAP visualization of scDisentangle’s progressive latent levels (levels 1, 3, 6, 8, and 16 shown) on Kang (top) and Hao (bottom) PBMCs, coloured by annotated cell type. UMAP was fit on the finest level (level 16) and applied to all levels. **b,** UMAP of the finest level coloured by condition (top; Kang) and by donor (bottom; Hao). **c,** Dendrograms constructed by iterative merging of cell-type groups across progressive latent levels on Kang (top) and Hao (bottom). **d,** Gene expression on NK cells across latent levels for Kang (top) and Hao (bottom). UMAPs are coloured by log-normalized expression (colour scale clipped at the 2nd and 98th percentiles) of *GZMB* (left), *PRF1* (middle), and by latent level (right). **e,** Matrixplot of mean log-normalized expression of *GZMB* (left) and *PRF1* (right) per latent level in NK cells on Kang (top) and Hao (bottom). Scaled between 0 and 1 across levels.

To quantify this ordering, we constructed dendrograms by stepping from fine to coarse latent levels and merging the two most similar groups at each step. Across both PBMC datasets, these dendrograms recapitulate canonical hematopoietic lineage relationships (Fig. 5c). On T cells from a B-cell acute lymphoblastic leukemia (B-ALL) cohort^26^, they largely capture T cell subset organization (Extended Data Fig. 9a). The principal lineage relationships were stable across random initializations (Extended Data Fig. 9b-d).

To examine how genes track this progressive structure, we used scDisentangle’s decoder to generate level-specific expression profiles. *GZMB* and *PRF1* expression increase on NK and CD8+ T cells (Fig. 5d,e; Supplementary Fig. 16) with increasing latent resolution. *GZMB* and *PRF1* are cytotoxic markers upregulated during activation and functional maturation of CD8+ T and NK cells^27^. Conversely, *MEF2C* is highest at the earliest levels and declines thereafter on T cells (Extended Data Fig. 10a), consistent with its role as a progenitor-associated transcription factor that is downregulated upon T cell maturation^28^. Additional markers show similar level-dependent patterns (Extended Data Fig. 10b-e).

Early progressive levels recovered broad expression structure shared across annotated immune populations, whereas later levels resolved finer cell-type and subtype distinctions. This extends to perturbation responses: on Kang, the shared IFN-β signature is captured at the earliest level while cell-type-specific effects appear at later levels (Supplementary Fig. 17).

## Discussion

Predicting cellular responses in held-out biological contexts is a central problem in single-cell biology. Despite recent progress, existing approaches struggle to outperform naive baseline average estimates^7,8^, and current evaluation frameworks do not assess whether predictions preserve the transcriptional identity of the source cell. scDisentangle generates counterfactuals that remain anchored to the individual source cell. This anchoring translated into more accurate predictions in held-out contexts and unseen perturbation combinations, and supported analyses at the level of individual donors and cell states, where population-level predictions remain limited.

On the introduced identity preservation metric, scDisentangle retained cell-specific structure that competing methods lost (Fig. 2c,g; Extended Data Figs. 4i and 6i). Separately, using a perturbed-centroid reference (Systema^8^), scDisentangle’s performance across the 128 Norman combinatorial perturbations was less correlated with systematic variation than that of CPA^3^ and GEARS^13^ (Fig. 2k). Intuitively, the two readouts probe different axes: one at the level of individual cells within a perturbation, the other across perturbations.

In immune checkpoint inhibitor–associated myocarditis^14^, we predicted how a given donor’s circulating CD8+ T cells would adapt upon heart infiltration in leave-one-donor-out configurations with no cardiac cells from that donor in training. Here, scDisentangle outperformed both the context-agnostic and the perturbation-agnostic baselines (Fig. 3b,c), indicating that neither a donor-invariant tissue effect nor an unchanged blood profile accounts for the observed adaptation. This result does not rely on the identity preservation score, providing independent evidence that predictions are donor-resolved.

Applying scDisentangle to a mouse prostate castration-regeneration time course^19^, we computed velocities between counterfactuals at consecutive held-out stages. Predicted *Ly6d* velocity rose at late castration and reversed upon androgen add-back (Extended Data Fig. 7e,f), consistent with the reported dynamics of this castration-resistant progenitor marker^24^. Cell-state transition probabilities also shifted across the cycle: L2-to-L1 transition probability declined progressively during castration and increased 5.4-fold at regeneration onset relative to the preceding transition (Fig. 4h), coinciding with re-entry of luminal cells into the cell cycle.

scDisentangle’s sequential decomposition also induces a progressive latent structure. Each unobserved factor is extracted from the residual left by preceding factors, encouraging successive factors to encode complementary variation. The resulting factors showed lower pairwise mutual information than those of CPA^3^ and scDisInFact^5^ (Extended Data Fig. 8c,d). Across two PBMC datasets^9,25^ and one B-ALL dataset^26^, dendrograms built by merging groups from fine to coarse latent levels recapitulated canonical hematopoietic lineage relationships and T cell subset organization, respectively (Fig. 5c; Extended Data Fig. 9a). The decoded level-specific profiles showed cytotoxic markers^27^ increasing with latent resolution, while a progenitor-associated transcription factor^28^ declined (Fig. 5d,e; Extended Data Fig. 10a). Early levels thus appear to encode a less differentiated transcriptional core that later levels refine into cell-type-specific programs. The same ordering applies to perturbation responses: on Kang, the shared IFN-β signature was captured at the earliest level, with less shared components at later levels (Supplementary Fig. 17). We regard this ordering as a descriptive property of the models we trained; whether it extends to tissues with more complex hierarchies remains open. Notably, on the two PBMCs, this was achieved without any progenitor populations in the training set, indicating that it reflects a property of mature transcriptional states rather than an observed differentiation trajectory.

Several limitations qualify the results we reported. In the liver, spatial zone and batch are not fully separable, and zone-specific effects should be read with this in mind. Prostate transition estimates rest on stages containing relatively few L2 cells, and RNA velocity estimates. Overall the predicted estimates are highly congruent with observed time course dynamics, but they are best read as a hypothesis about luminal progenitor expansion timing. Furthermore, single-cell RNA sequencing is destructive, making it presently impossible to observe truly matched ground truth for validating per-cell counterfactuals. Hence, predictions are validated at the population level, using correlation, distributional, and RMSE-type metrics. Finally, scDisentangle requires each attribute value to be observed in training and does not use structural priors over perturbations. It therefore predicts responses in unseen contexts and unseen combinations of observed perturbations, but not responses to perturbations absent from training, as knowledge-graph and chemical-structure-based methods attempt^13,29^.

Further directions follow. scDisentangle treats observed attributes as discrete labels; extending attribute contributions to continuous covariates might allow dose and time to be modelled directly, but is not straightforward using the current formulation. Conversely, the step-wise discretisation of the latent space, while preserving the continuous space of individual neurons, offers straightforward interpretability and could be used as an effective surrogate where needed (e.g. in cases like lineage tracing). The level decomposition also suggests a way to ask at which level of biological resolution a perturbation acts, e.g. whether a compound shifts the shared transcriptional core or only cell-type-specific programs, providing deeper informative insights than single predicted profiles. Finally, we suggest that source-cell identity preservation be reported alongside population-level accuracy in perturbation benchmarks, as faithful representation of the perturbed cell identity and perturbation prediction may diverge. A method that performs well on the latter alone may not effectively capture true biological heterogeneity and support the cell- and donor-resolved analyses that motivate counterfactual prediction.

Virtual-cell models will ultimately be judged by whether they can predict how a given cellular state responds to perturbations and their combinations in previously unobserved contexts. The scale, sparsity and nonlinearity of these response spaces place that goal beyond immediate reach. scDisentangle advances a tractable part of this problem by composing predictions from observed perturbations while preserving the source-cell identity on which those perturbations act. More broadly, our results suggest that forward progress will require improved benchmarks, and models that retain biological heterogeneity while disentangling separable biological components.

## Methods

### scDisentangle model

Consider a dataset 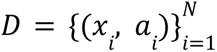 where 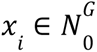 is the gene expression count vector for cell i across *G* genes and 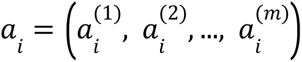 denote m discrete labelled attributes (e.g. cell type, genetic perturbation). scDisentangle jointly learns three components: an encoder-decoder (*E*_θ_, *G*_φ_) that learns a prior latent representation *d z_i_* ∈ *R^d^* from log-transformed counts; a decomposition module that factorizes *z_i_* into attribute-associated and unobserved factors; and a composition module that maps these factors back to *R^d^* for latent recovery and counterfactual generation. The encoder computes *z_i_* = *E*_θ_ (*log*(1 + *x_i_*)). The decoder *G*_φ_ parameterizes a per-gene Negative Binomial (NB) likelihood conditioned on *z_i_* and the cell’s log-library size, with learnable gene-wise inverse dispersion parameters.

### Perturbation prediction

scDisentangle’s decomposition separates cell-intrinsic variation 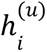 from attribute-related effects. Predicting the transcriptional response of a control cell to perturbation *p* thus reduces to setting the target attribute to *p* and re-running the composition module with 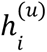 held fixed, then decoding the resulting composed latent. At inference, when individual components of a combinatorial perturbation are observed in the training set, scDisentangle calibrates gene-wise prediction bias using the average difference between predicted and observed single-perturbation responses.

### Evaluation metric

Predictions were evaluated using correlation (*R*², Pearson, *R*² delta, Pearson delta), error (RMSE, Normalized RMSE), and distributional (Wasserstein) metrics, computed on top-K differentially expressed genes per condition. Normalized RMSE divides the RMSE between predicted and observed mean profiles by the RMSE between the observed perturbed and control means. Additional combinatorial-perturbation-specific metrics were computed for Norman benchmarks. Negative metric values were set to 0 in the text, figures and significance tests. Supplementary Tables report uncapped values.

### Identity preservation score

For each predicted cell, we rank the Euclidean distance to its source control cell (matched by cell identifier) among distances to all control cells and define the preservation score as 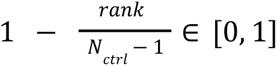, averaged over cells.

### Competing methods

We compared scDisentangle with CPA^3^, scGen^1^, scDisInFact^5^, biolord^6^ and GEARS^13^. For perturbation-prediction benchmarks, all models were trained following their respective published tutorials with recommended hyperparameters.

## Data availability

The Kang^9^ dataset is available from the Gene Expression Omnibus under accession GSE96583. The preprocessed version used here was obtained from trVAE^30^ repository: (https://drive.google.com/drive/folders/1n1SLbXha4OH7j7zZ0zZAxrj_-2kczgl8). The Liver expression matrix was downloaded from figshare: (https://figshare.com/articles/dataset/spatio-temporal-infection_infected/22148900?file=39375713), as provided by biolord^6^. The perturb-seq dataset^11^ was downloaded from Zenodo (https://zenodo.org/records/7041849; scPerturb^31^). We used the Gene Expression Omnibus GSE228597 and GSE146811 accession numbers to obtain the Myocarditis^14^ and the Prostate^19^ datasets, respectively. We obtained the B-ALL^26^ dataset from CELLxGENE (https://cellxgene.cziscience.com/collections/14dc301f-d4fb-4743-a590-aa88d5f1df1a), and the Hao et al.^25^ dataset via scvi-tools^32^ (v0.20.3; `scvi.data.pbmc_seurat_v4_cite_seq`).

## Supporting information

Supplementary Table 2 - Kang benchmark

Supplementary Table 3 - Liver benchmark

Supplementary Table 5 - Split IDs of double perturbations in the Norman combinatorially-seen scenario

Supplementary Table 6 - Benchmark on Norman combinatorially-seen scenario using standard DEGs

Supplementary Table 7 - Benchmark on Norman combinatorially-seen scenario using combination-specific DEGs

Supplementary Table 9 - Benchmark on Norman single-only scenario using standard DEGs

Supplementary Table 10 - Benchmark on Norman single-only scenario using combination-specific DEGs

Supplementary Table 11 - Benchmark on myocarditis CD8+ T subset

Supplementary Table 12 - Benchmark on myocarditis using the full dataset

## Acknowledgements

We thank the authors of the original studies for making the datasets used in this work publicly available.

## Author Contributions

S.E.M. and S.E. led the conceptualization of the study, with R.E.F., M.H. and T.D. S.E.M. and S.E. implemented the method, performed the data analysis, and prepared the figures. A.B. contributed to the preparation of the figures and to the revision of the manuscript. F.Z.E.H. contributed to the design of the evaluation metrics and to the biological interpretation of the myocarditis analysis. R.E.F., M.H. and T.D. guided the design of the method and the analyses, contributed to the interpretation of the results, and supervised the study. S.E.M. and S.E. wrote the manuscript with input from R.E.F., M.H. and T.D. All authors reviewed and approved the final manuscript.

## Ethics

All datasets used in this study are publicly available from published studies with ethics oversight in the original studies. No new human or animal experiments were performed.

## Extended Data

**Extended Data Fig. 1:**
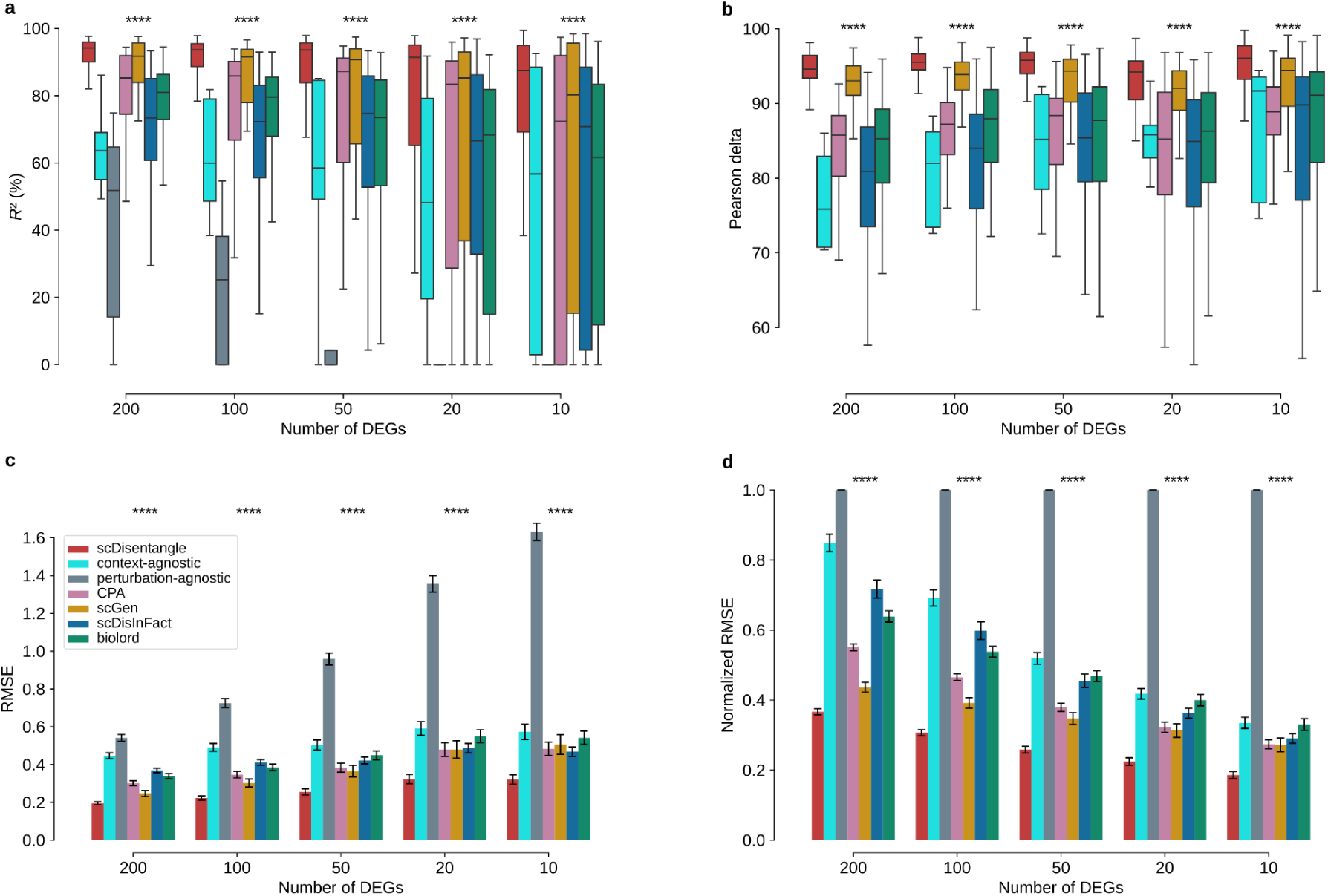
Benchmark on Kang using additional metrics. **a-d,** Benchmark of scDisentangle against the context-agnostic and perturbation-agnostic baselines, CPA, scGen, scDisInFact and biolord on *R*² (**a**; coefficient of determination), Pearson delta (**b**; Pearson correlation computed on the perturbation-induced change relative to control; perturbation-agnostic baseline excluded as it serves as the reference for delta computation), RMSE (**c**; root mean squared error), and normalized RMSE (**d**) across *n* = 80 evaluations (8 cell types x 10 seeds) on Kang. In boxplots (**a**, **b**), the centre line indicates median, box edges the first and third quartiles, and whiskers extend to 1.5× the interquartile range; outliers not shown. Bars (**c**,**d**) indicate the mean and error bars the standard error of the mean (s.e.m.). Significance was assessed using one-sided Mann-Whitney U tests applied within each OOD context against the best-performing competing method at each DEG count; context-level p-values were combined using Fisher’s method; \**P* < 0.05, \*\**P* < 0.01, \*\*\**P* < 0.001, \*\*\*\**P* < 0.0001.

**Extended Data Fig. 2:**
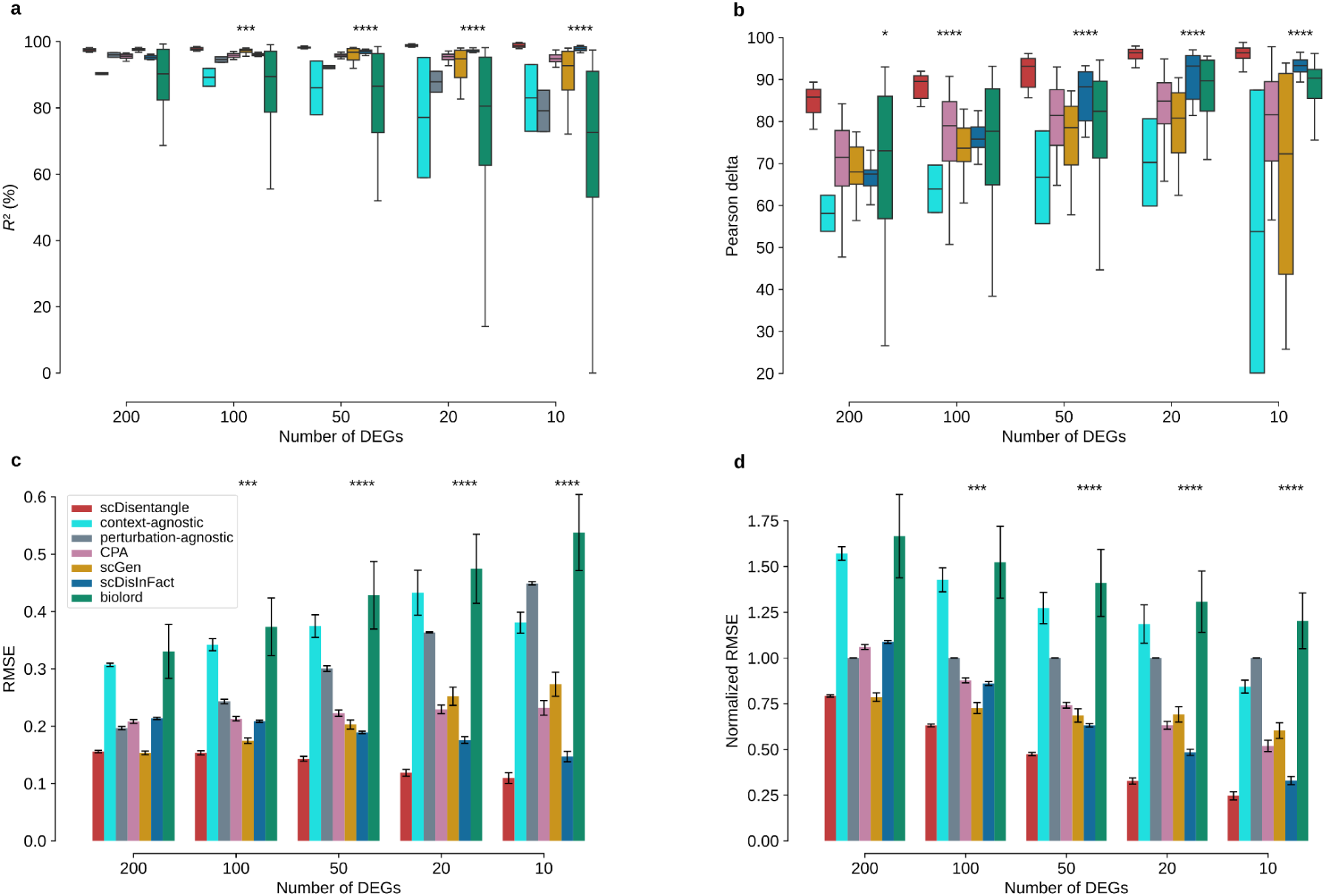
Benchmark on Liver using additional metrics. **a-d,** Benchmark of scDisentangle against naive baselines, CPA, scGen, scDisInFact and biolord on *R*² (**a**), Pearson delta (**b**; perturbation-agnostic baseline excluded as it serves as the reference for delta computation), RMSE (**c**), and normalized RMSE (**d**) across *n* = 20 evaluations (2 zones x 10 random seeds) on Liver. In boxplots (**a**, **b**), the centre line indicates median, box edges the first and third quartiles, and whiskers extend to 1.5× the interquartile range; outliers not shown. Bars (**c**,**d**) indicate the mean and error bars the standard error of the mean (s.e.m.). Significance was assessed using one-sided Mann-Whitney U tests applied within each OOD context against the best-performing competing method at each DEG count; context-level p-values were combined using Fisher’s method; \**P* < 0.05, \*\**P* < 0.01, \*\*\**P* < 0.001, \*\*\*\**P* < 0.0001.

**Extended Data Fig. 3:**
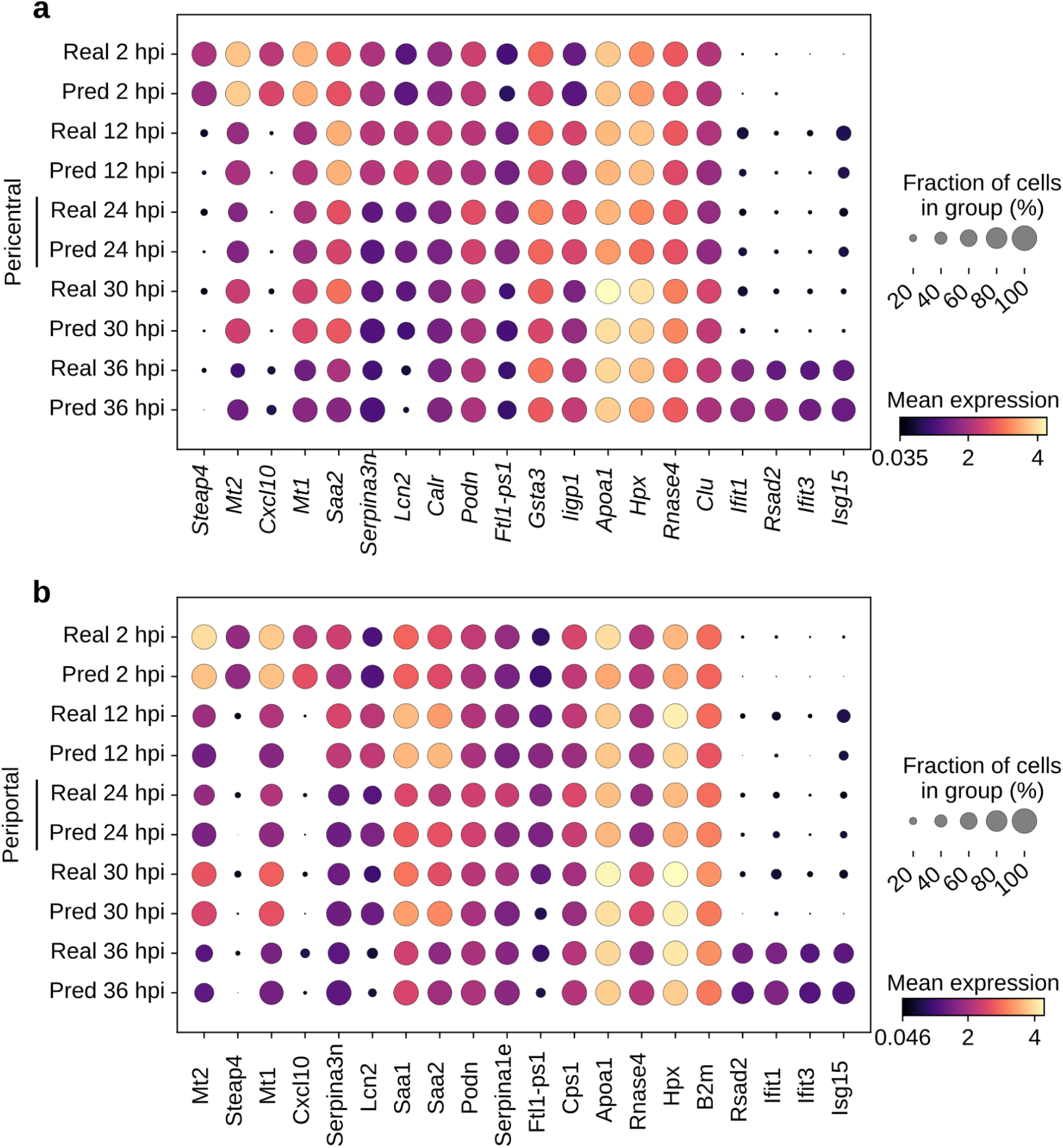
Comparing observed and predicted expressions across liver infection time-course. **a,b,** Dot plot comparing observed vs. scDisentangle-predicted log-normalized expression of infection time-specific marker genes across the infection time course on cells of the pericentral zone (**a**; reproduced from Fig. 2j for comparison) and periportal zone (**b**).

**Extended Data Fig. 4:**
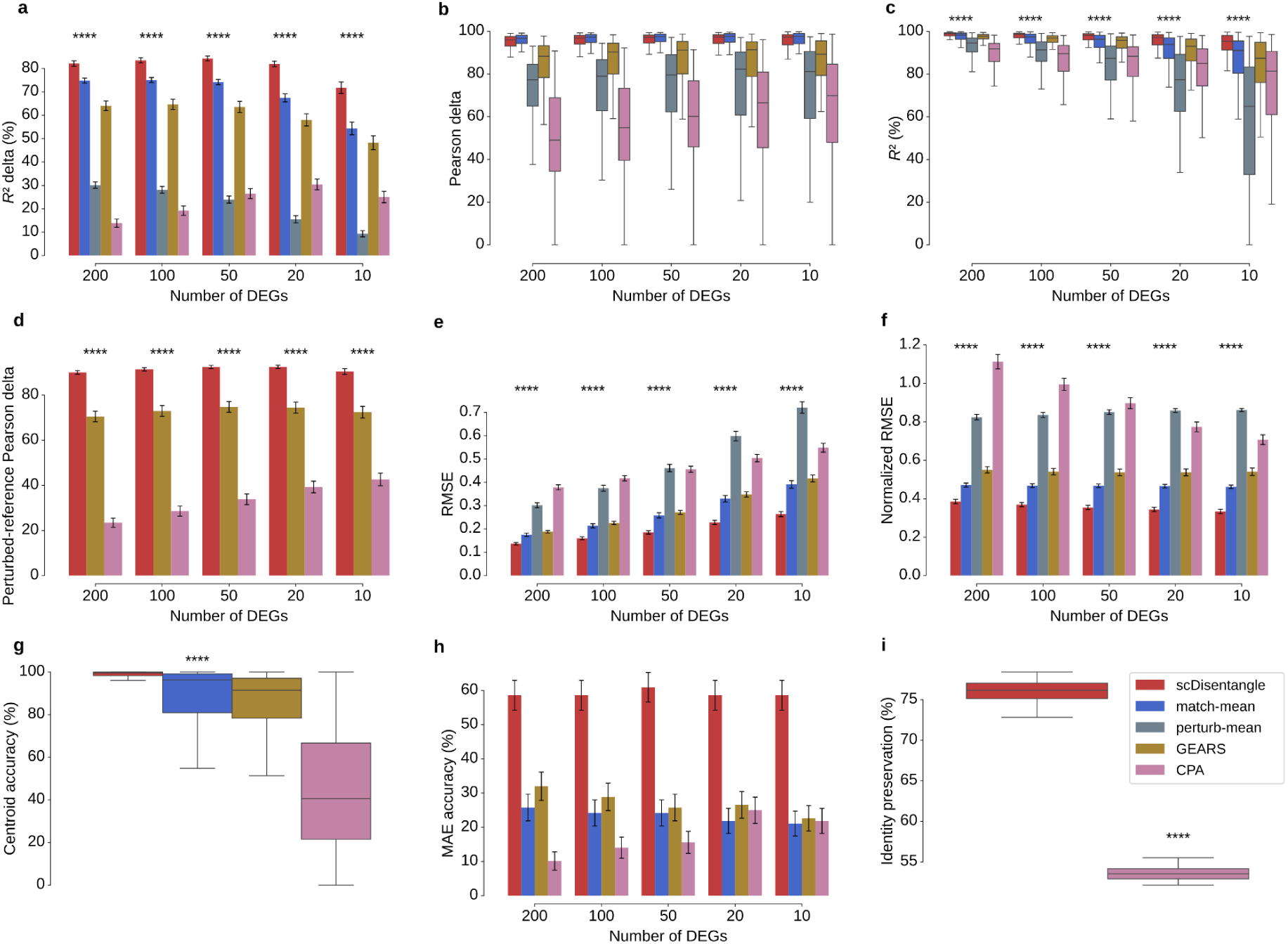
Benchmark on Norman combinatorially-seen scenario using standard DEGs. **a-i,** Benchmark of scDisentangle against the match-mean and perturb-mean baselines where applicable, GEARS and CPA on *R*² delta (**a**), Pearson delta (**b**), *R*² (**c**), perturbed-reference Pearson delta (**d**), RMSE (**e**), normalized RMSE (**f**), centroid accuracy (**g**), MAE (mean absolute error) accuracy (**h**), and identity preservation (**i**) across *n* = 128 evaluations (corresponding to 128 double-perturbations) on Norman (combinatorially-seen out-of-distribution scenario; perturbation-vs-control DEGs). Bars (**a**, **d**, **e**, **f**, **h**) indicate the mean and error bars the standard error of the mean (s.e.m.). In boxplots (**b**, **c**, **g**, **i**), the centre line indicates median, box edges the first and third quartiles, and whiskers extend to 1.5× the interquartile range; outliers not shown. Significance was assessed using a one-sided paired Wilcoxon signed-rank test against the best-performing competing method at each DEG count. \**P* < 0.05, \*\**P* < 0.01, \*\*\**P* < 0.001, \*\*\*\**P* < 0.0001. Significance stars are only shown when the median difference favours scDisentangle.

**Extended Data Fig. 5:**
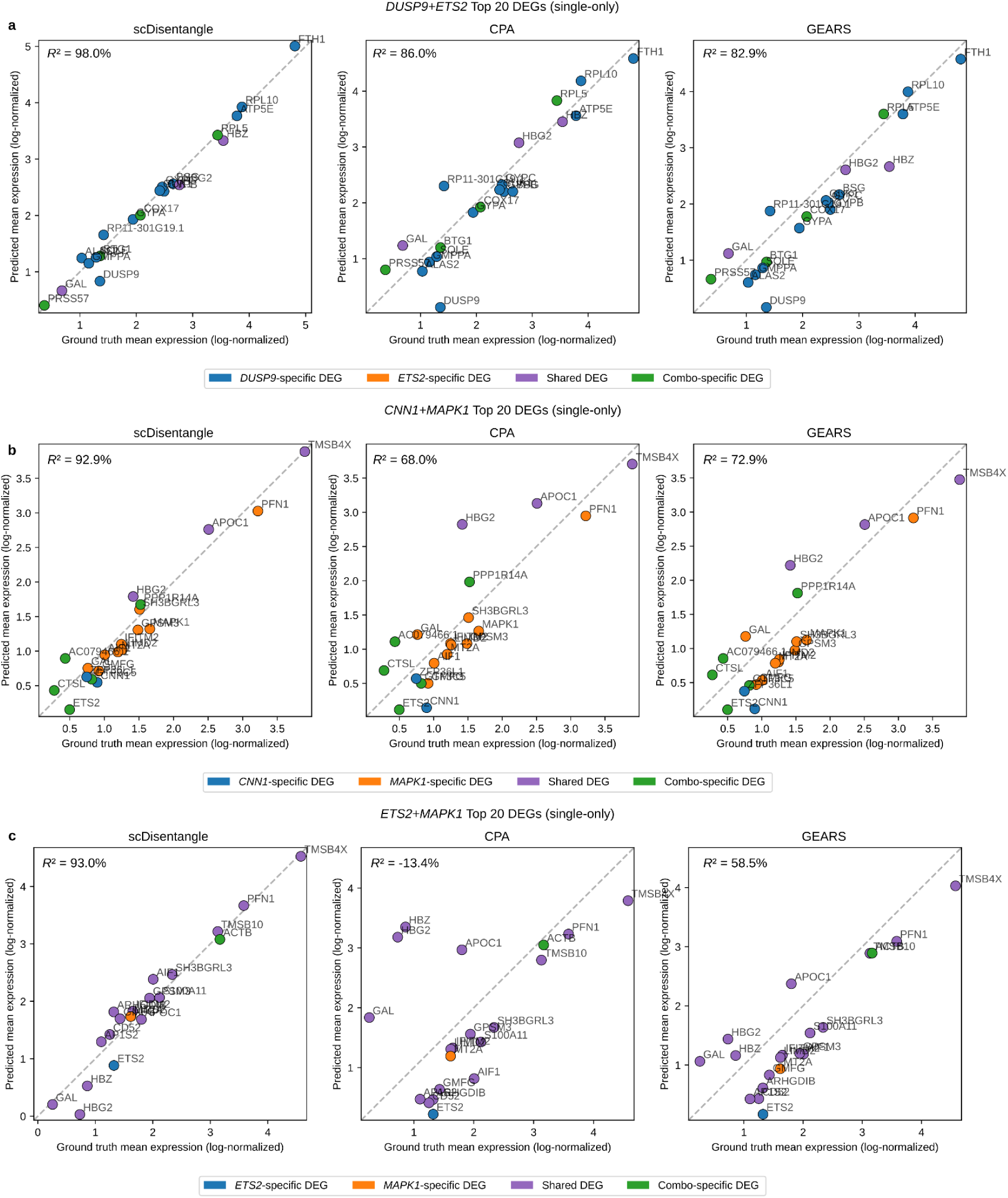
Comparing observed and predicted expressions on Norman single-only scenario. **a-c,** Predicted vs. observed mean log-normalized expression for *DUSP9*+*ETS2* (**a**), *CNN1*+*MAPK1* (**b**), and *ETS2*+*MAPK1* (**c**) combinatorial perturbations (single-only out-of-distribution scenario); layout, gene colouring, and statistics as in Fig. 2l. Neither constituent is seen in combination during training.

**Extended Data Fig. 6:**
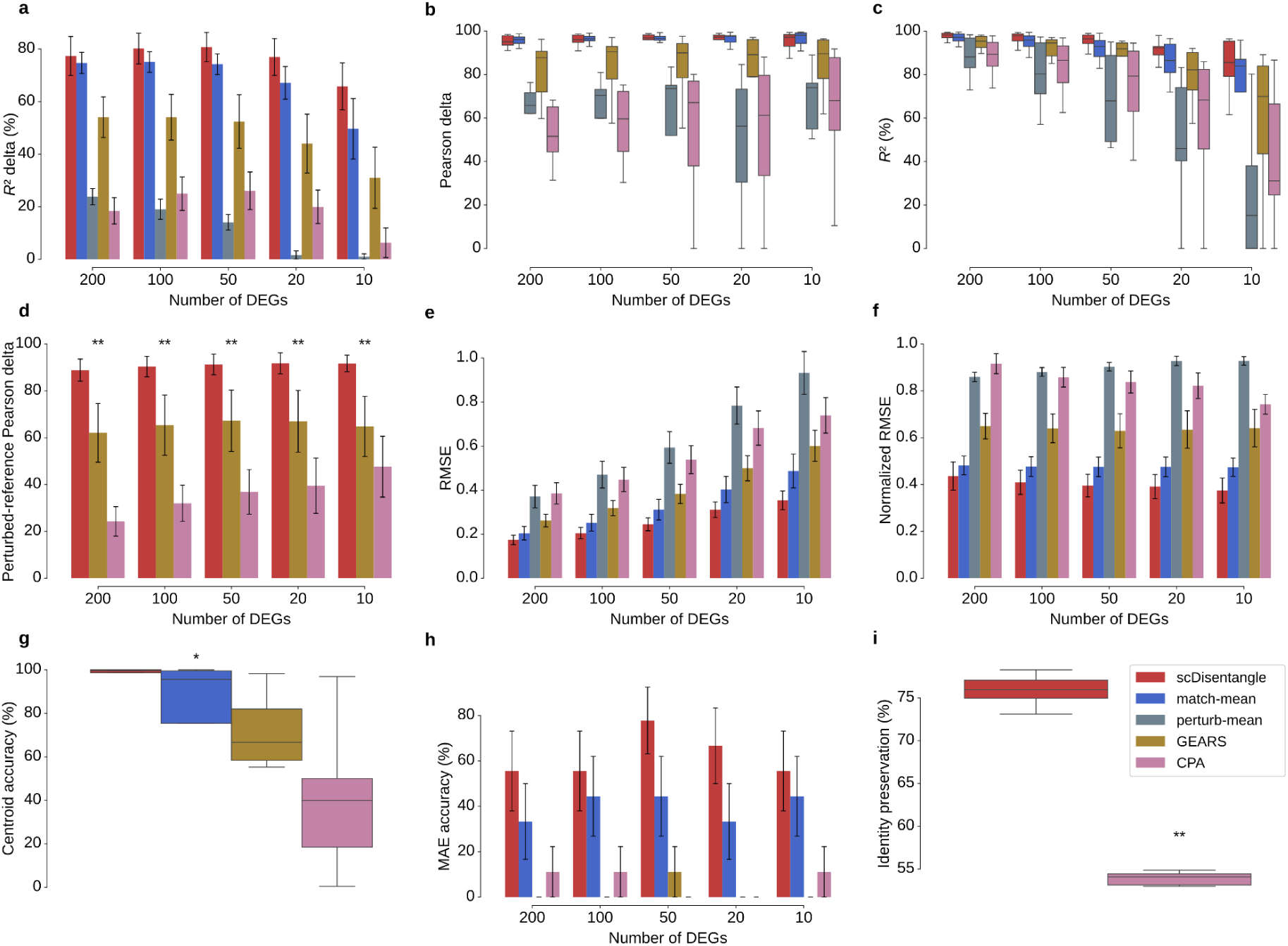
Benchmark on Norman single-only scenario using standard DEGs. **a-i,** Benchmark of scDisentangle against the match-mean and perturb-mean baselines where applicable, GEARS and CPA on *R*² delta (**a**), Pearson delta (**b**), *R*² (**c**), perturbed-reference Pearson delta (**d**), RMSE (**e**), normalized RMSE (**f**), centroid accuracy (**g**), MAE (mean absolute error) accuracy (**h**) and identity preservation (**i**) across *n* = 9 evaluations on Norman (single-only out-of-distribution scenario; perturbation-vs-control DEGs). Bars (**a**, **d**, **e**, **f**, **h**) indicate the mean and error bars the standard error of the mean (s.e.m.). In boxplots (**b**, **c**, **g**, **i**), the centre line indicates median, box edges the first and third quartiles, and whiskers extend to 1.5× the interquartile range; outliers not shown. Significance was assessed using a one-sided paired Wilcoxon signed-rank test against the best-performing competing method at each DEG count; \**P* < 0.05, \*\**P* < 0.01, \*\*\**P* < 0.001, \*\*\*\**P* < 0.0001. Significance stars are only shown when the median difference favours scDisentangle.

**Extended Data Fig. 7:**
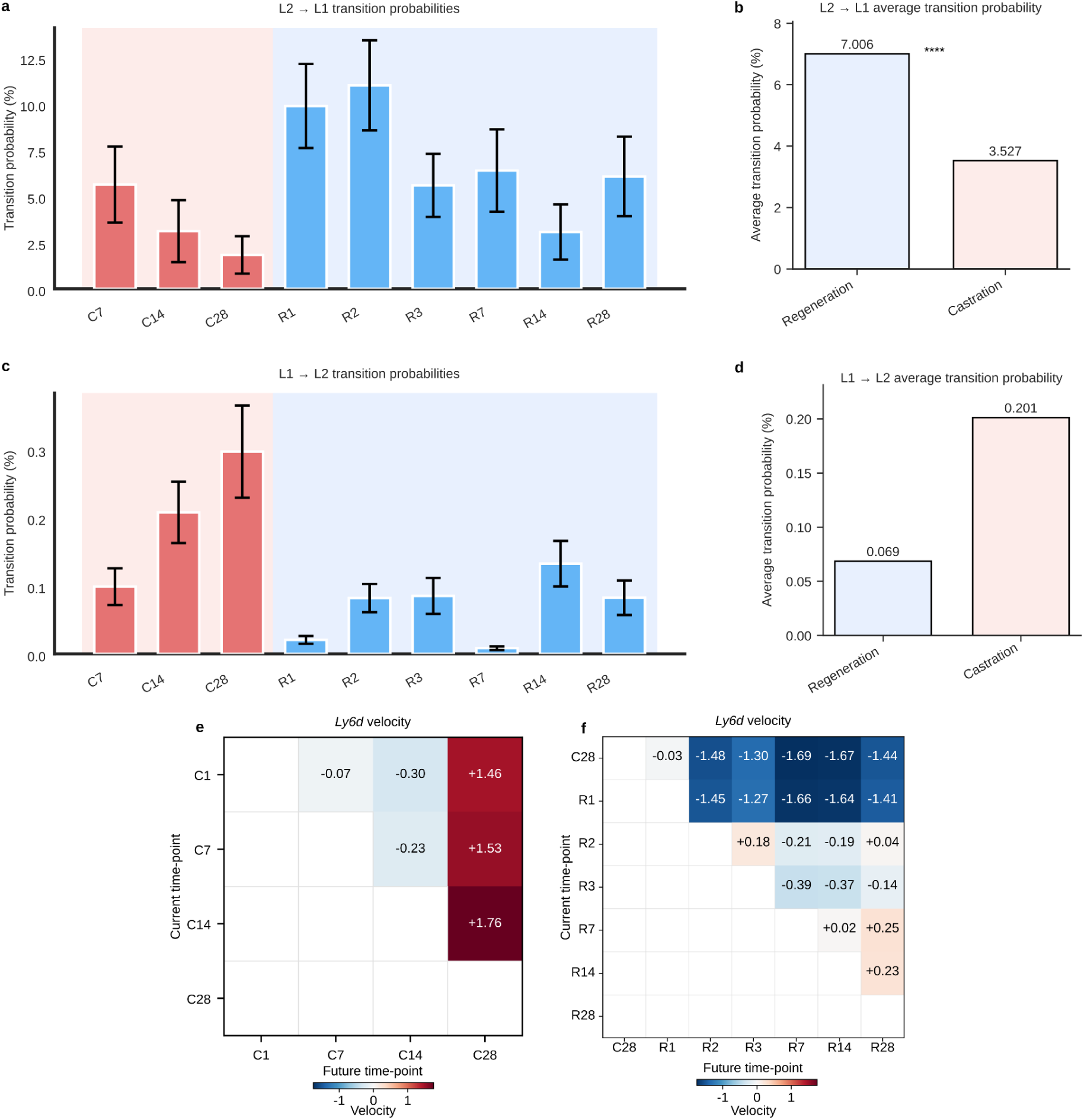
Prostate cell-cell transition analysis. **a,** Per-transition L2-to-L1 transition probabilities (%) for each pair of consecutive scDisentangle-generated C/R stages. Bars indicate the mean and error bars the s.e.m across *n* = 97 L2 cells. Self-transitions excluded. **b,** Mean L2-to-L1 transition probability averaged over castration and regeneration stages, as in Fig. 4i, reproduced here for direct comparison with panel **d**. **c,** Per-transition L1-to-L2 transition probabilities (%), as in (**a**) across *n* = 3,211 L1 cells. **d,** Mean L1-to-L2 transition probability averaged over castration and regeneration stages. Statistical significance assessed using one-sided Wilcoxon signed-rank test (**b**, regeneration > castration; **d**, castration > regeneration); \**P* < 0.05, \*\**P* < 0.01, \*\*\**P* < 0.001, \*\*\*\**P* < 0.0001. **e,** Mean scDisentangle-generated *Ly6d* velocity (future - current expression) for L2 cells across castration stages. Rows: current state; columns: future state. Reproduced from Fig. 4j for comparison. **f,** Mean scDisentangle-generated *Ly6d* velocity in L2 cells across regeneration stages (from C28 onward). Rows: current state; columns: future state.

**Extended Data Fig. 8:**
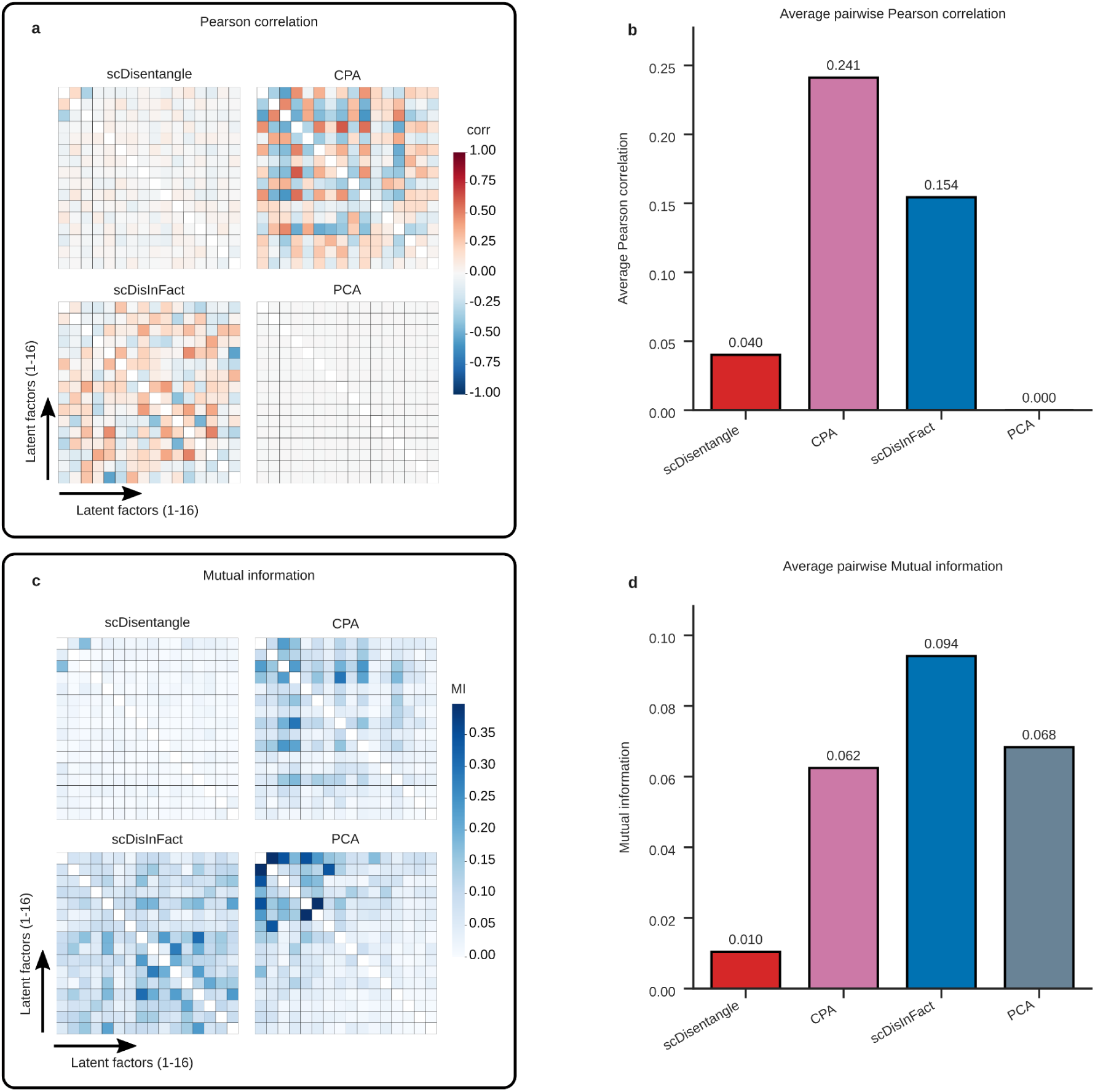
Pairwise dependence between latent dimensions. **a,** Heatmap of pairwise Pearson correlation between latent dimensions of scDisentangle, CPA, scDisInFact, and PCA on Kang. Colour scale shared across methods. **b,** Average absolute pairwise Pearson correlation. **c,** Heatmap of pairwise mutual information between latent dimensions of scDisentangle, CPA, scDisInFact, and PCA on Kang. Colour scale shared across methods and capped at the 99th percentile. **d,** Average pairwise mutual information. Latent dimensionality was matched across methods: CPA was trained with n_latent = 16, scDisInFact with a 16-dimensional shared biological latent, and PCA was restricted to its first 16 principal components.

**Extended Data Fig. 9:**
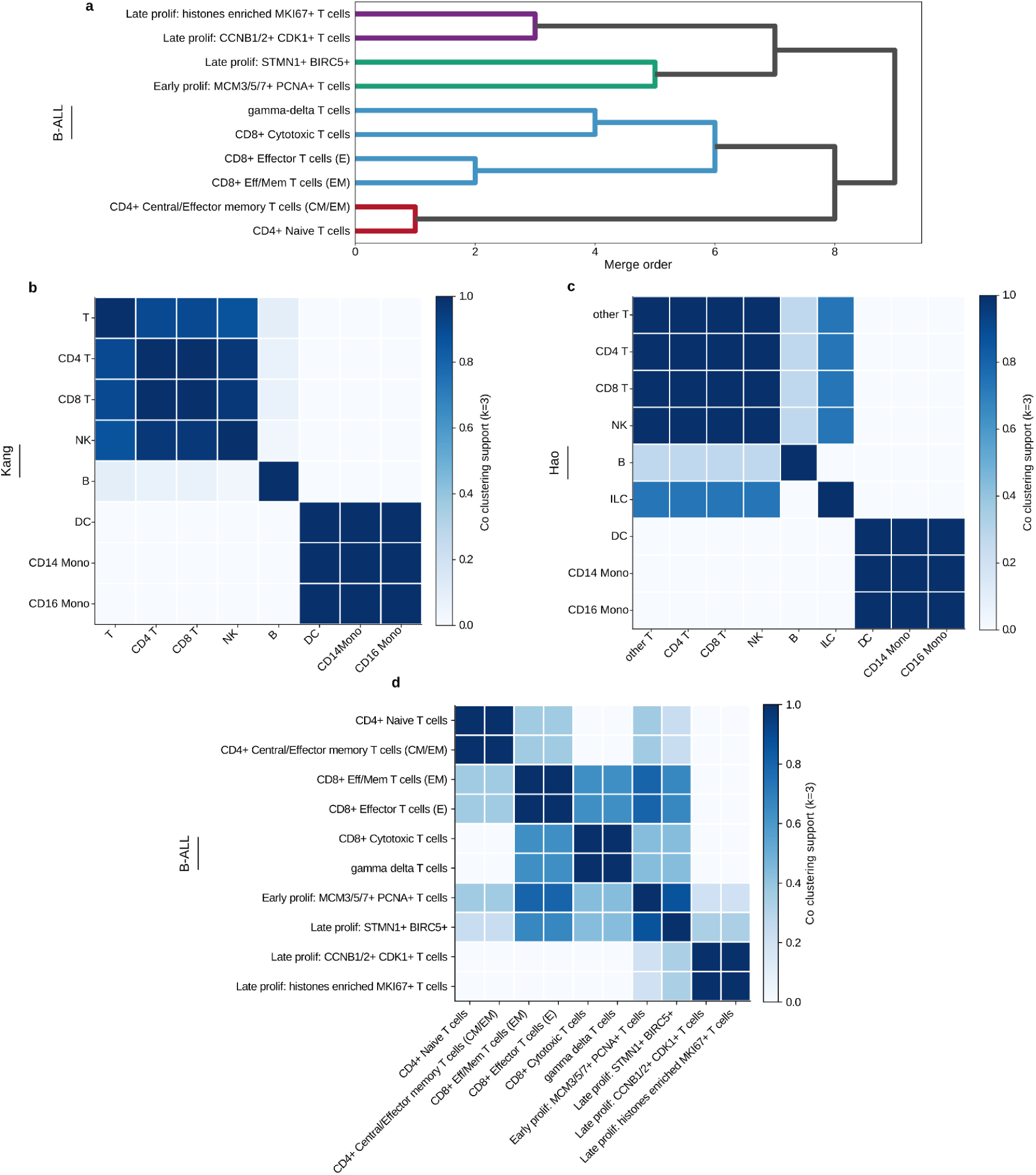
Dendrograms inferred from scDisentangle progressive latents. **a**, Dendrogram constructed by iterative merging of cell-type groups across progressive latent levels on B-ALL dataset. **b-d,** Co-clustering support heatmap for Kang (**b**), Hao (**c**), and B-ALL (**d**). The dendrogram construction algorithm was run 30 times with different random seeds (varying GMM initializations). For each run, the resulting dendrogram was cut into k = 3 groups. Each entry shows the fraction of runs in which a given pair of cell types was assigned to the same cluster.

**Extended Data Fig. 10:**
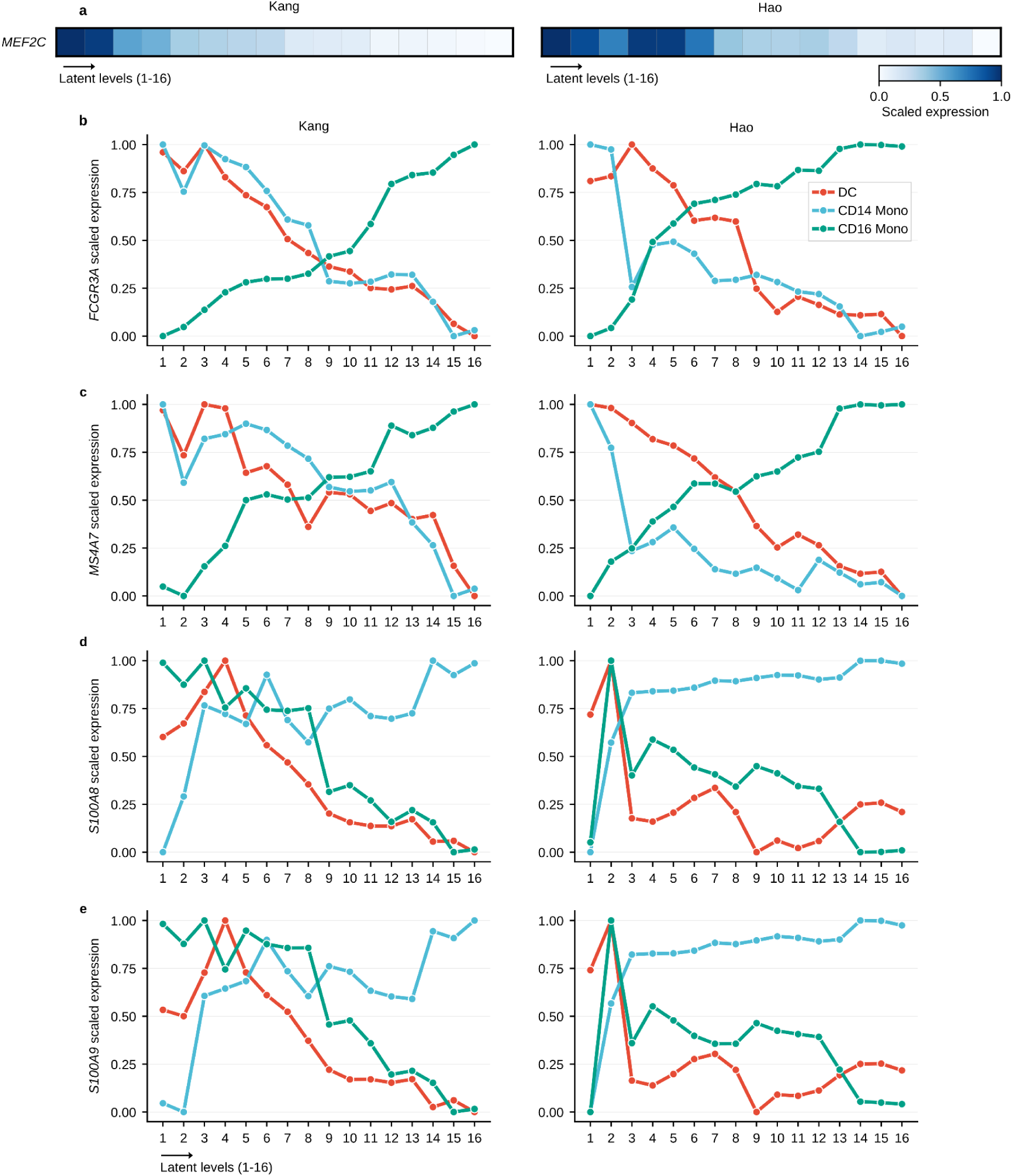
Gene expression patterns across decoded progressive latents. **a,** Matrixplot of mean log-normalized expression of *MEF2C* per latent level in T cells on Kang (left) and Hao (right). Scaled between 0 and 1 across levels. **b-e,** Line plots of scaled mean log-normalized expression of *FCGR3A* (**b**), *MS4A7* (**c**), *S100A8* (**d**), and *S100A9* (**e**), on dendritic cells (DC; red), CD14+ monocytes (light blue), and CD16+ monocytes (green), on Kang (left) and Hao (right). Values are scaled between 0 and 1 per cell type across levels.

