## Supplementary Information (Supplementary Figures 1-19, Supplementary Tables 1-17) for "scDisentangle: sequential disentanglement for single-cell perturbation response prediction"

**Supplementary Tables** 4
Supplementary Table 1: Dataset summary. 4
Supplementary Table 2: Kang benchmark. 4
Supplementary Table 3: Liver benchmark. 4
Supplementary Table 4: Perturbations excluded from the Norman dataset during preprocessing. 5
Supplementary Table 5: Split IDs of double perturbations in the Norman combinatorially-seen scenario. 5
Supplementary Table 6: Benchmark on Norman combinatorially-seen scenario using standard DEGs. 5
Supplementary Table 7: Benchmark on Norman combinatorially-seen scenario using combination-specific DEGs. 5
Supplementary Table 8: Out-of-distribution perturbations and their genetic interaction group assignments in the Norman single-only scenario. 6
Supplementary Table 9: Benchmark on Norman single-only scenario using standard DEGs. 6
Supplementary Table 10: Benchmark on Norman single-only scenario using combination-specific DEGs. 6
Supplementary Table 11: Benchmark on myocarditis CD8+ T subset. 7
Supplementary Table 12: Benchmark on myocarditis using the full dataset. 7
Supplementary Table 13: Hyperparameters used for scDisentangle training. 8
Supplementary Table 14: CD8+ T cell annotation markers. 8
Supplementary Table 15: Genes retained during preprocessing of the prostate dataset.10
Supplementary Table 16: Cell populations removed from the Hao et al. PBMC dataset during preprocessing. 10
Supplementary Table 17: Training computational cost as a function of batch size. 11
**Supplementary Figures** 12
Supplementary Fig. 1: Comparison of real and predicted gene expression distributions. 13
Supplementary Fig. 2: Benchmark on Norman combinatorially-seen scenario using combination-specific DEGs. 14
Supplementary Fig. 3: Performance of models as a function of systematic variation. 15
Supplementary Fig. 4: Comparison of predicted and observed expressions in Norman single-only scenario. 16
Supplementary Fig. 5: Comparison of predicted and observed expressions in Norman single-only scenario on two additional combinatorial perturbations. 17
Supplementary Fig. 6: Benchmark on Norman single-only scenario using combination-specific DEGs. 18
Supplementary Fig. 7: Benchmark on Norman using distributional metrics. 19
Supplementary Fig. 8: Benchmark on Norman using Perturbed-reference $R^2$ delta. 20
Supplementary Fig. 9: Benchmark against naive baselines on Myocarditis (CD8+ T subset). 21
Supplementary Fig. 10: Benchmark against naive baselines on Myocarditis (full data). 22
Supplementary Fig. 11: Co-clustering and divergence of luminal L1 and L2 cells. 23
Supplementary Fig. 12: Comparing average observed and scDisentangle-generated expression across the castration-regeneration cycle. 24
Supplementary Fig. 13: Comparing observed and scDisentangle-generated expression

for additional genes across the castration-regeneration cycle. 25
Supplementary Fig. 14: Visualization of scDisentangle's progressive latents on Kang PBMCs. 26
Supplementary Fig. 15: Visualization of scDisentangle's progressive latents on Hao PBMCs. 27
Supplementary Fig. 16: Gene expression patterns across scDisentangle's generated levels. 28
Supplementary Fig. 17: Perturbation effect decomposition analysis on Kang. 29
Supplementary Fig. 18: Perturbation identity preservation for combinatorial perturbations (combinatorially-seen scenario). 30
Supplementary Fig. 19: Benchmark on Liver spatio-temporal experiment. 31
**Supplementary References** **32**

### Supplementary Tables

| Dataset | Organism | Perturbation | Cells | Genes | Groups |
|---|---|---|---|---|---|
| Kang et al.[1] | Human | IFN-β stimulation | 13,576 | 13,404 | 8 (cell types) |
| Afriat et al.[2] | Mouse | Plasmodium infection | 19,053 | 8,203 | 2 (zones) |
| Norman et al.[3] | Human (K562) | CRISPRa | 108,619 | 5,446 | 1 (cell-line) |
| Blum et al.[4] | Human | Tissue | 32,573 | 5,000 | 12 (donors) |
| Karthaus et al.[5] | Mouse | C/R cycle time-points | 87,187 | 5,027 | 25 (cell types) |
| Hao et al.[6] | Human | – | 93,542 | 5,000 | 9 (cell types) |
| Guerrero et al.[7] | Human | – | 37,100 | 5,000 | 11 (cell types) |

**Supplementary Table 1: Dataset summary.**

For each dataset, the organism, perturbation type, number of cells, number of genes and number of biological groups are reported.

**Supplementary Table 2: Kang benchmark.**

Results using top-20 DEGs, averaged per cell type (10 seeds per cell type). Identity preservation is computed on the full gene set. Values are reported uncapped. (Provided as a separate Excel file.)

**Supplementary Table 3: Liver benchmark.**

Results using top-20 DEGs, averaged per zone (10 seeds per zone). Identity preservation is computed on the full gene set. Values are reported uncapped. (Provided as a separate Excel file.)

| Perturbation | Type | Reason for removal |
|---|---|---|
| C19orf26 | Single | Absent from expression matrix |
| C3orf72 | Single | Absent from expression matrix |
| IER5L | Single | Unsupported by GEARS GO graph |
| KIAA1804 | Single | Absent from expression matrix |
| C3orf72+FOXL2 | Double | Contains C3orf72 |
| LYL1+IER5L | Double | Unsupported by GEARS GO graph |
| TGFBR2+C19orf26 | Double | Contains C19orf26 |

**Supplementary Table 4: Perturbations excluded from the Norman dataset during preprocessing.**

Each perturbation is listed alongside its type (single or double) and reason for removal.

**Supplementary Table 5: Split IDs of double perturbations in the Norman combinatorially-seen scenario.**

The split ID indicates the split from which each perturbation was held out during training. (Provided as a separate Excel file.)

**Supplementary Table 6: Benchmark on Norman combinatorially-seen scenario using standard DEGs.**

Results are reported for each OOD perturbation (128 total). Metrics are computed on top-20 DEGs (OOD perturbation vs. control). For centroid accuracy and identity preservation, the full gene set is used. Values are reported uncapped. (Provided as a separate Excel file.)

**Supplementary Table 7: Benchmark on Norman combinatorially-seen scenario using combination-specific DEGs.**

Results are reported for each OOD perturbation (128 total). Metrics are computed on top-20 DEGs (OOD perturbation vs. single constituents). For centroid accuracy and identity preservation, the full gene set is used, as in Supplementary Table 6. Values are reported uncapped. (Provided as a separate Excel file.)

| Perturbation | Genetic interaction |
|---|---|
| CBL+CNN1 | STRONG SYNERGY SIMILAR PHENOTYPES |
| DUSP9+ETS2 | EPISTASIS |
| DUSP9+MAPK1 | NEOMORPHIC |
| ETS2+MAPK1 | REDUNDANT |
| ETS2+CEBPE | -- |
| CNN1+MAPK1 | SUPPRESSORS |
| CEBPB+CEBPA | REDUNDANT |
| CBL+PTPN12 | STRONG SYNERGY SIMILAR PHENOTYPES |
| AHR+FEV | STRONG SYNERGY DISSIMILAR PHENOTYPES |

**Supplementary Table 8: Out-of-distribution perturbations and their genetic interaction group assignments in the Norman single-only scenario.**

Each perturbation is listed alongside its genetic interaction group, as identified by Norman et al.

**Supplementary Table 9: Benchmark on Norman single-only scenario using standard DEGs.**

Results are reported for each OOD perturbation (9 perturbations total). Metrics are computed on top-20 DEGs (OOD perturbation vs. control). For centroid accuracy and identity preservation, the full gene set is used. Values are reported uncapped. (Provided as a separate Excel file.)

**Supplementary Table 10: Benchmark on Norman single-only scenario using combination-specific DEGs.**

Results are reported for each OOD perturbation (9 perturbations total). Metrics are computed on top-20 DEGs (OOD perturbation vs. single constituents). For centroid accuracy and identity preservation, the full gene set is used, as in Supplementary Table 9. Values are reported uncapped. (Provided as a separate Excel file.)

**Supplementary Table 11: Benchmark on myocarditis CD8+ T subset.**

Results are averaged per OOD donor (11 donors; 10 seeds per donor). Metrics are computed on top-20 DEGs. Values are reported uncapped. (Provided as a separate Excel file.)

**Supplementary Table 12: Benchmark on myocarditis using the full dataset.**

Results are averaged per OOD donor (11 donors; 10 seeds per donor). Metrics are computed on top-20 DEGs. Values are reported uncapped. (Provided as a separate Excel file.)

| **Hyperparameter** | **Perturbation prediction** | **Latent structure analysis** |
|---|---|---|
| Latent factors | 16 | 16 |
| Base encoder | [n_genes, 256, 16] | [n_genes, 256, 16] |
| Sequential encoders | [16, 256, 1] | [16, 256, 1] |
| Sequential mappers | [d_a+t, 256, 16] (t = 1..16) | [d_a+t, 256, 16] (t = 1..16) |
| Activation (all MLPs) | Sinusoidal (Sine) | Sinusoidal (Sine) |
| Weight initialization | Xavier | Xavier |
| Batch normalization | None | None |
| DecoderSCVI layers | 3 | 3 |
| DecoderSCVI hidden size | 1,024 | 256 |
| DecoderSCVI batch/layer norm | None | None |
| Likelihood | Negative Binomial | Negative Binomial |
| Optimizer | AdamW | AdamW |
| Weight decay | 0.01 | 0.01 |
| Gradient clipping (max norm) | 1.0 | 1.0 |
| Initial sequential encoder + Base Auto-encoder learning rate | $1 \times 10^{-4}$ | $1 \times 10^{-4}$ |
| Residual encoders and mappers learning rate | $1 \times 10^{-3}$ | $1 \times 10^{-3}$ |
| Embedders learning rate | $1 \times 10^{-3}$ | $1 \times 10^{-3}$ |
| px_r learning rate | $1 \times 10^{-3}$ | $1 \times 10^{-3}$ |
| λ_rec (reconstruction loss weight) | $1 \times 10^{-5}$ | $1 \times 10^{-5}$ |

| | | |
|---|---|---|
| λ_latent (latent recovery loss weight) | $2 \times 10^{-3}$ | $2 \times 10^{-3}$ |
| Batch size | 128 | 128 |
| Training epochs | 521 | 521 |
| Checkpoint criterion | $R^2$ Mean (top 10 DEGs) | Mutual Information Gap (binned) |

**Supplementary Table 13: Hyperparameters used for scDisentangle training.**

Hyperparameter values are reported separately for the two tasks: perturbation prediction and latent structure analysis. d_a is the total attribute-embedding dimension. Mapper t takes the concatenation of the attribute embeddings and the first t unobserved factors.

| Score | Gene |
|---|---|
| cd4_score | IL7R |
| cd4_score | CCR7 |
| cd4_score | LST1 |
| cd4_score | SELL |
| cd8_score | CD8A |
| cd8_score | CD8B |
| cd8_score | GZMH |
| cd8_score | PRF1 |
| cd8_score | NKG7 |
| nk_score | NCAM1 |
| nk_score | KLRD1 |
| nk_score | KLRB1 |
| nk_score | FCGR3A |
| nk_score | NKG7 |

**Supplementary Table 14: CD8+ T cell annotation markers.**

Marker genes used for CD8+ T cell annotation in the myocarditis dataset.

| Gene | Category |
|---|---|
| Ly6a | Luminal progenitor marker |
| Psca | Luminal progenitor marker |
| Tacstd2 | Luminal progenitor marker |
| Nkx3-1 | Luminal marker |
| Pbsn | Luminal secretory marker |
| Dpp4 | L1 luminal marker |
| Prom1 | L1 luminal marker |
| Cd59a | Surface marker |
| Egf | Growth factor |
| Trp63 | Basal marker |
| Krt5 | Basal keratin |
| Krt14 | Basal keratin |
| Cd24a | Surface marker |
| Krt8 | Luminal keratin |
| Krt18 | Luminal keratin |
| Foxi1 | Ionocyte marker |
| Nrg2 | Growth factor |
| Rspo3 | Wnt signalling |
| Krt13 | Keratin |
| Wnt2 | Wnt ligand |
| Wnt6 | Wnt ligand |
| Wnt10a | Wnt ligand |
| Rorb | Transcription factor |
| Rspo1 | Wnt signalling |
| Fgf10 | Growth factor |
| Sult1e1 | Estrogen metabolism |
| Acta2 | Smooth muscle marker |
| Myh11 | Smooth muscle marker |
| Notch3 | Notch signalling |
| Fgfr2 | Growth factor receptor |
| Igf1 | Growth factor |
| Lgr4 | Wnt signalling |

| | |
|---|---|
| Mki67 | Proliferation marker |
| Cldn* family (35 genes) | Claudin tight junction family |

**Supplementary Table 15: Genes retained during preprocessing of the prostate dataset.**

Each gene is listed alongside its category. These genes were selected based on the original study of Karthaus et al. and were explicitly preserved prior to highly variable gene selection.

| Removed population | Annotation level | Category |
|---|---|---|
| HSPC | celltype.l1 | Precursor |
| Platelet | celltype.l1 | Non-immune |
| Eryth | celltype.l1 | Erythroid |
| CD4 Naive | celltype.l3 | Naive |
| CD8 Naive | celltype.l3 | Naive |
| CD8 Naive_2 | celltype.l3 | Naive |
| CD4 TCM_1 | celltype.l3 | Central memory |
| CD4 TCM_2 | celltype.l3 | Central memory |
| CD4 TCM_3 | celltype.l3 | Central memory |
| CD8 TCM_1 | celltype.l3 | Central memory |
| CD8 TCM_2 | celltype.l3 | Central memory |
| CD8 TCM_3 | celltype.l3 | Central memory |
| B naive kappa | celltype.l3 | Naive |
| B naive lambda | celltype.l3 | Naive |
| B intermediate kappa | celltype.l3 | Intermediate |
| B intermediate lambda | celltype.l3 | Intermediate |
| NK_CD56bright | celltype.l3 | CD56bright NK |
| NK Proliferating | celltype.l3 | Proliferating |
| CD4 Proliferating | celltype.l3 | Proliferating |
| CD8 Proliferating | celltype.l3 | Proliferating |
| ASDC_mDC | celltype.l3 | AS dendritic cells |
| ASDC_pDC | celltype.l3 | AS dendritic cells |

**Supplementary Table 16: Cell populations removed from the Hao et al. PBMC dataset during preprocessing.**

Each population is listed alongside its annotation level and category.

| Batch Size | Time/Epoch (sec) | Peak GPU Memory (GB) | Est. Full Training (min) |
|---|---|---|---|
| 128 | 3.14 | 3.28 | 27.2 |
| 256 | 1.91 | 3.34 | 16.6 |
| 512 | 1.27 | 3.45 | 11.0 |
| 1024 | 0.99 | 3.73 | 8.6 |

**Supplementary Table 17: Training computational cost as a function of batch size.**

Benchmarks were performed on the Kang PBMC dataset (11,521 training cells from 13,576 total x 13,404 genes) using a single NVIDIA RTX A6000 GPU. Estimated full training assumes 521 epochs.

### Supplementary Figures

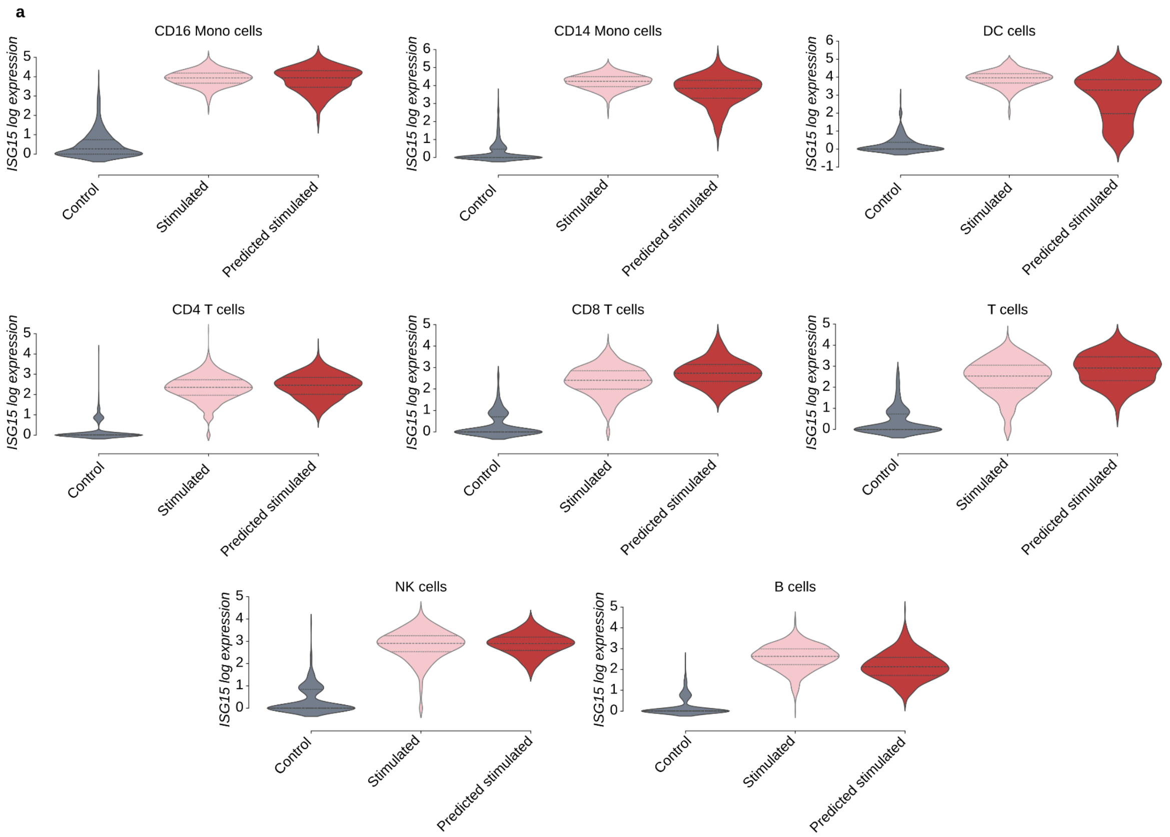


**Supplementary Fig. 1: Comparison of real and predicted gene expression distributions.**

**a,** Violin plots comparing the log-normalized expression distributions of real control, real stimulated, and scDisentangle-predicted stimulated cells for the top differentially expressed gene (stimulated vs. control) in each annotated Kang cell type (seed = 1). Horizontal lines indicate the median and quartiles.

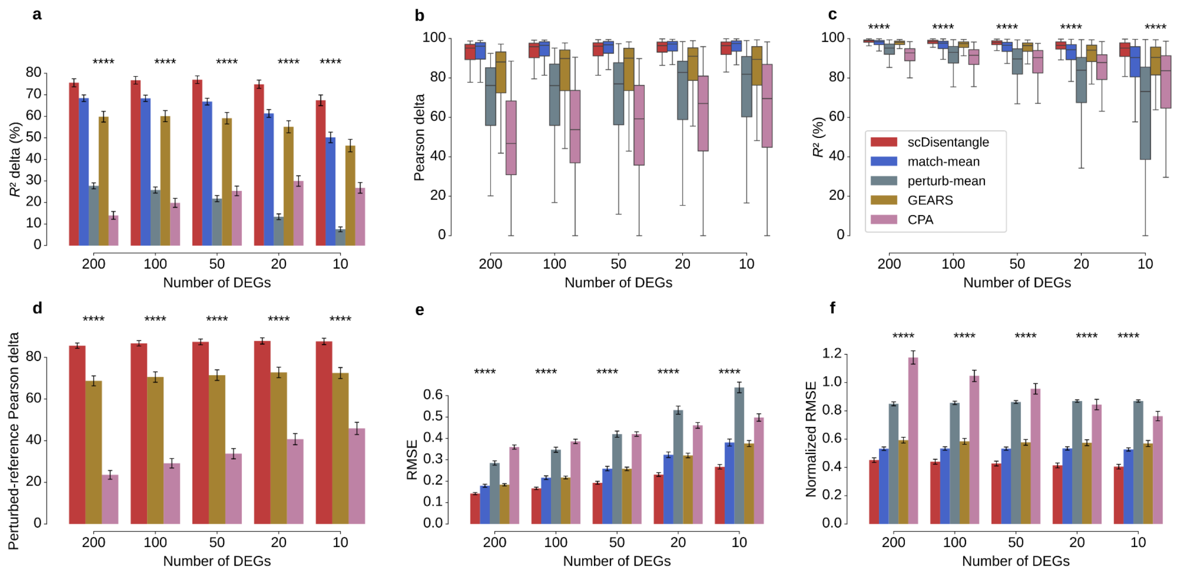


**Supplementary Fig. 2: Benchmark on Norman combinatorially-seen scenario using combination-specific DEGs.**

**a-f,** Benchmark of scDisentangle against the match-mean and perturb-mean baselines where applicable, GEARS and CPA on $R^2$ delta (**a**), Pearson delta (**b**), $R^2$ (**c**), perturbed-reference Pearson delta (**d**), RMSE (**e**), and Normalized RMSE (**f**) across $n$ = 128 evaluations (corresponding to 128 double-perturbations) on Norman (combinatorially-seen OOD scenario; perturbation-vs-single-constituents DEGs). Bars (**a**, **d**, **e**, **f**) indicate the mean and error bars the standard error of the mean (s.e.m.). In boxplots (**b**, **c**), centre line indicates median, box edges the first and third quartiles, and whiskers extend to 1.5× the interquartile range; outliers not shown. Significance was assessed using a one-sided paired Wilcoxon signed-rank test against the best-performing competing method at each DEG count. *$P$ < 0.05, **$P$ < 0.01, ***$P$ < 0.001, ****$P$ < 0.0001.

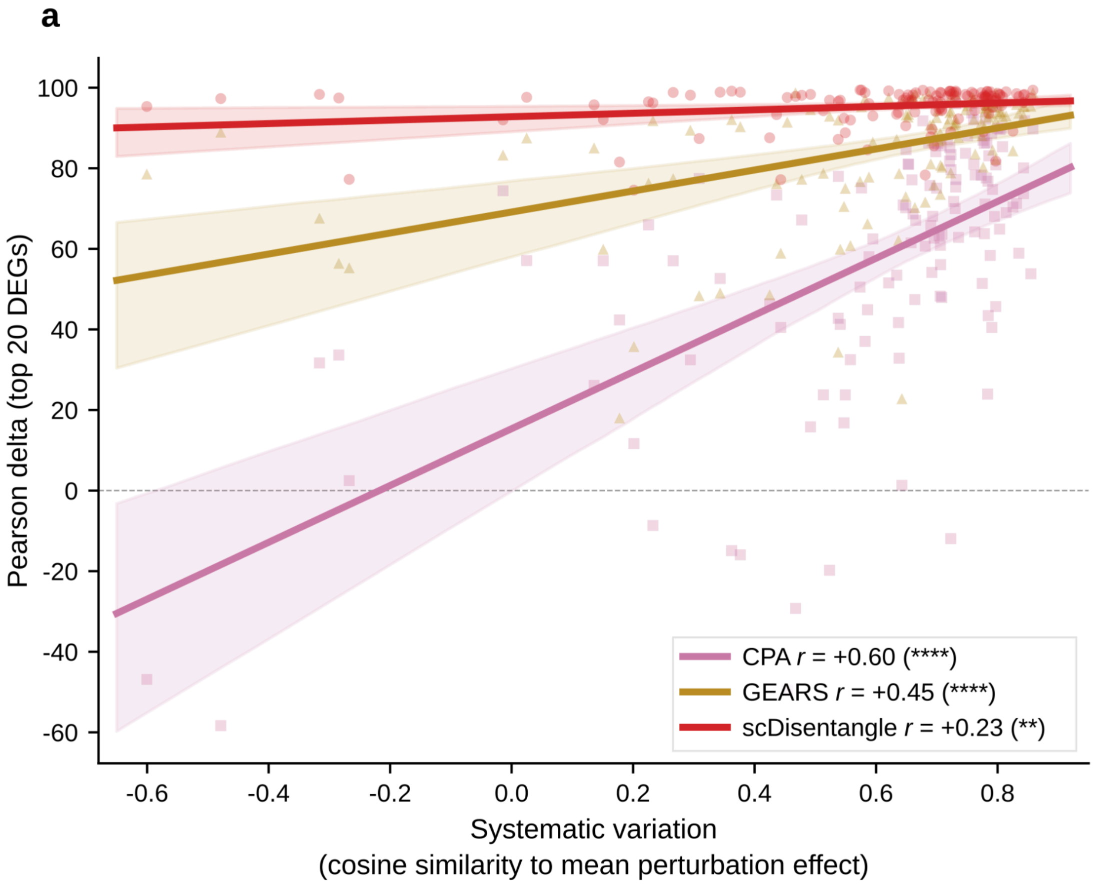


**Supplementary Fig. 3: Performance of models as a function of systematic variation.**

**a,** For each of 128 combinatorial perturbations in Norman, systematic variation is quantified as the cosine similarity between the perturbation's differential expression profile (perturbation centroid - control centroid) and the mean perturbation effect (centroid of all perturbed cells - control centroid). The y-axis shows each model's Pearson delta (the Pearson correlation between predicted and observed expression profiles after subtracting the mean control profile) evaluated on the top 20 DEGs. Lines show linear regression fits; *r* denotes the Pearson correlation between systematic variation and model performance across perturbations, with significance assessed by a two-sided Pearson correlation test; n.s., not significant, *$P < 0.05$, **$P < 0.01$, ***$P < 0.001$, ****$P < 0.0001$. Shaded bands show bootstrapped 95% confidence intervals (2,000 resamples).

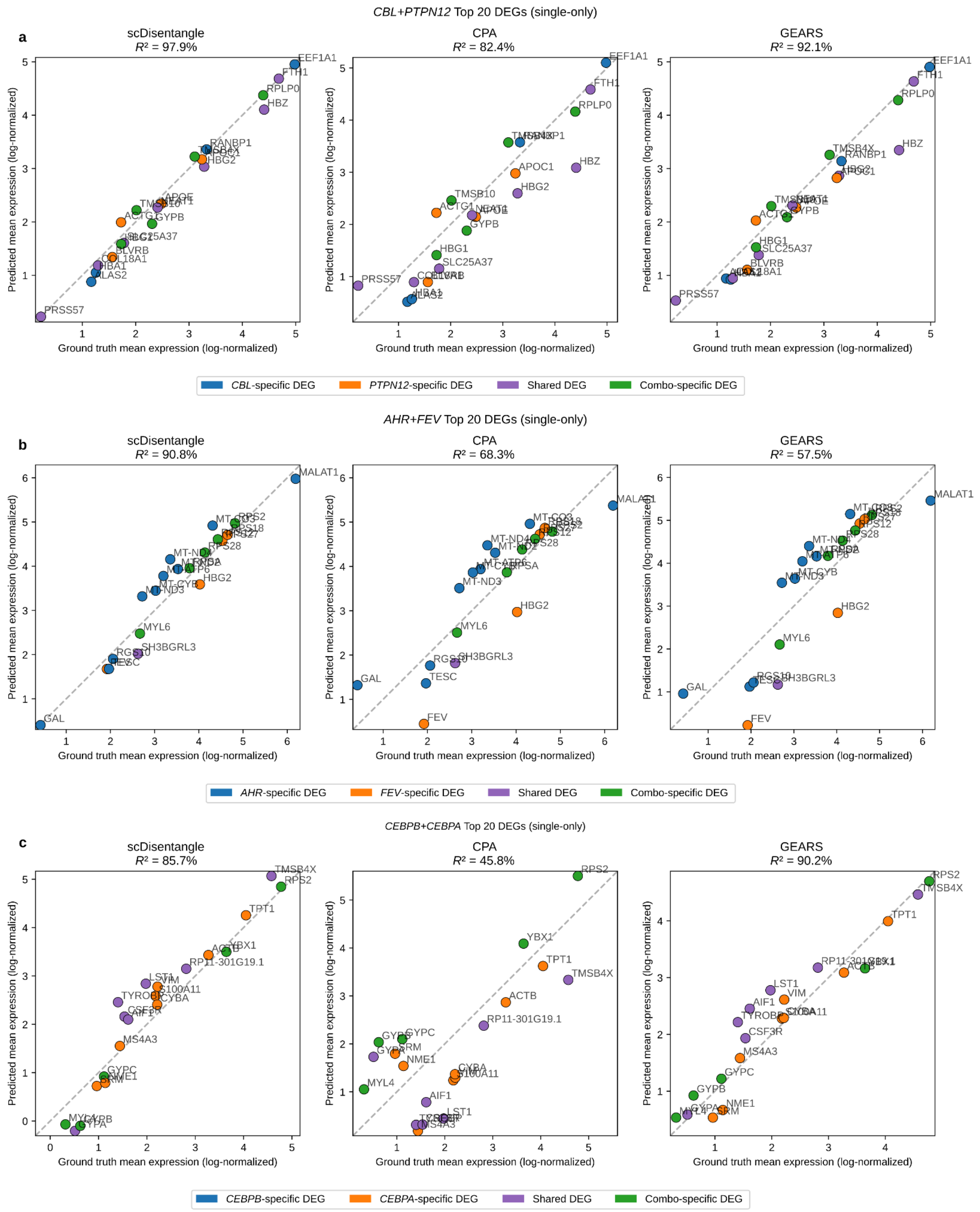


**Supplementary Fig. 4: Comparison of predicted and observed expressions in Norman single-only scenario.**

**a-c,** Predicted vs. observed mean log-normalized expression for *CBL+PTPN12* (**a**), *AHR+FEV* (**b**), and *CEBPB+CEBPA* (**c**) combinatorial perturbations (single-only OOD scenario); layout, gene colouring, and statistics as in Fig. 2l.

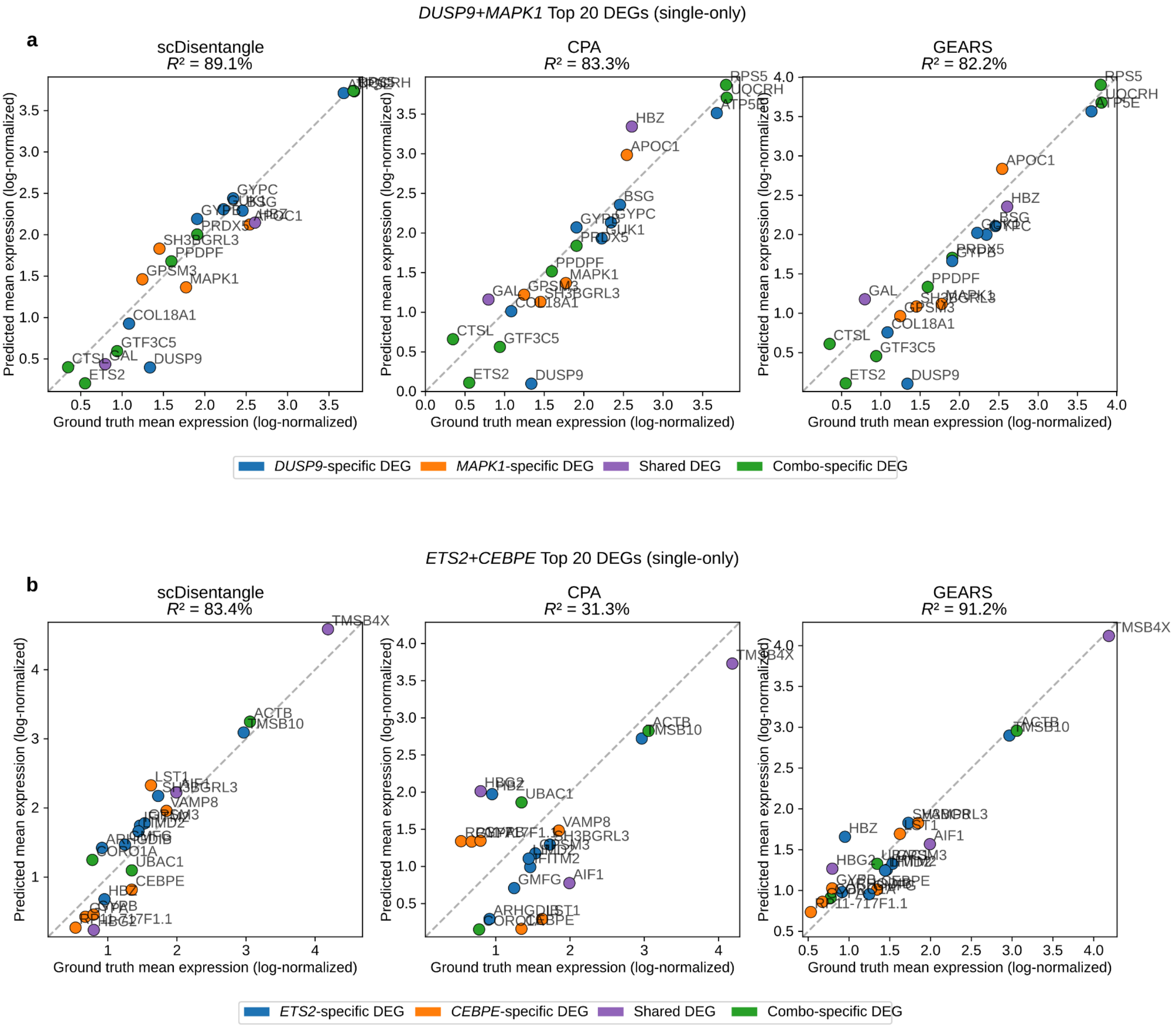


**Supplementary Fig. 5: Comparison of predicted and observed expressions in Norman single-only scenario on two additional combinatorial perturbations.**

**a,b,** Predicted vs. observed mean log-normalized expression for *DUSP9+MAPK1* (**a**) and *ETS2+CEBPE* (**b**) combinatorial perturbations (single-only OOD scenario); layout, gene colouring, and statistics as in Fig. 2l.

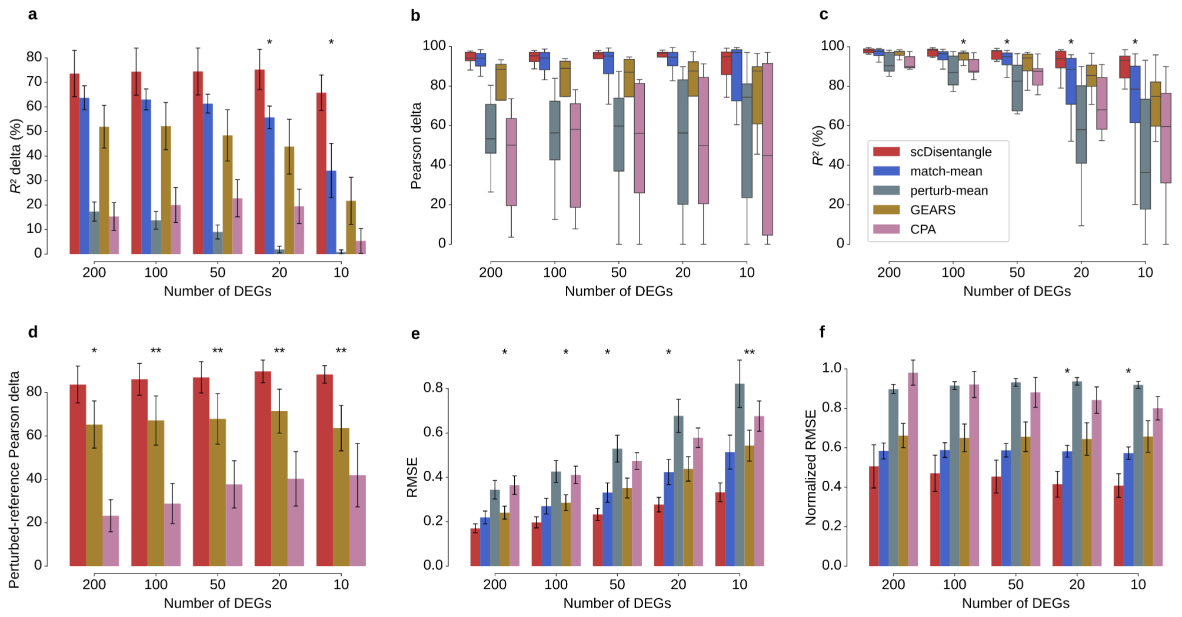


**Supplementary Fig. 6: Benchmark on Norman single-only scenario using combination-specific DEGs.**

**a-f,** Benchmark of scDisentangle against the match-mean and perturb-mean baselines where applicable, GEARS and CPA on $R^2$ delta (**a**), Pearson delta (**b**), $R^2$ (**c**), perturbed-reference Pearson delta (**d**), RMSE (**e**) and Normalized RMSE (**f**) across $n$ = 9 evaluations (corresponding to nine OOD perturbations) on Norman (single-only OOD scenario; perturbation-vs-single-constituents DEGs). Bars (**a**, **d**, **e**, **f**) indicate the mean and error bars the standard error of the mean (s.e.m.). In boxplots (**b**, **c**), centre line indicates median, box edges the first and third quartiles, and whiskers extend to 1.5× the interquartile range; outliers not shown. Significance was assessed using a one-sided paired Wilcoxon signed-rank test against the best-performing competing method at each DEG count. *$P$ < 0.05, **$P$ < 0.01, ***$P$ < 0.001, ****$P$ < 0.0001.

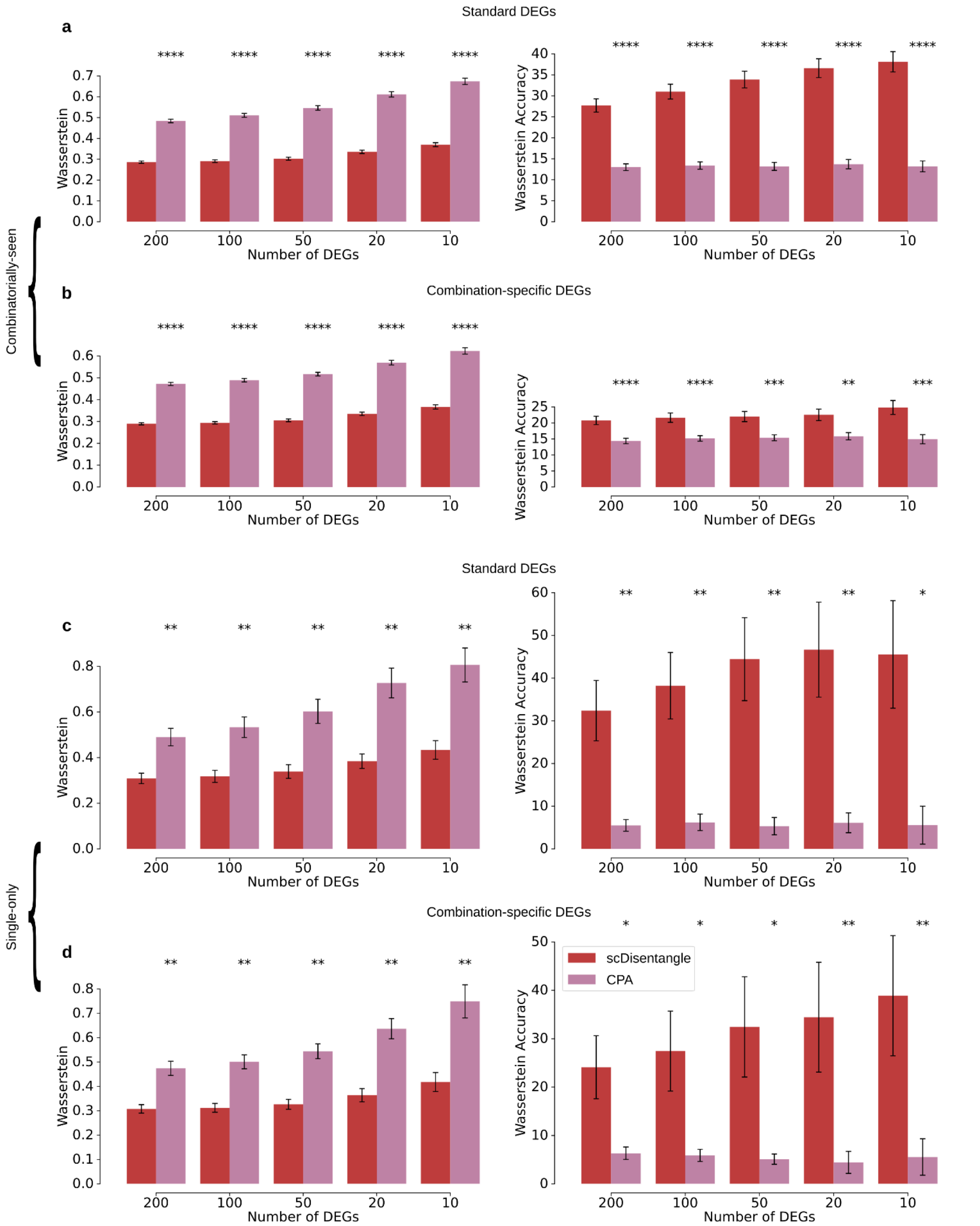


**Supplementary Fig. 7: Benchmark on Norman using distributional metrics.**
**a-d,** Wasserstein (left) and Wasserstein accuracy (right) evaluated on standard DEGs (**a**, **c**) and combination-specific DEGs (**b**, **d**) across $n$ = 128 evaluations (combinatorially-seen OOD scenario; **a**, **b**) and $n$ = 9 evaluations (single-only OOD scenario; **c**, **d**) on Norman. Wasserstein accuracy is the proportion of genes for which the predicted distribution is closer to the true double-perturbation distribution than to the control or either single-constituent distribution. Bars indicate the mean and error bars the s.e.m. Significance was assessed using a one-sided paired Wilcoxon signed-rank test against CPA at each DEG count; *$P$ < 0.05, **$P$ < 0.01, ***$P$ < 0.001, ****$P$ < 0.0001. GEARS and naive baselines are excluded since they output a mean effect vector rather than per-cell response.

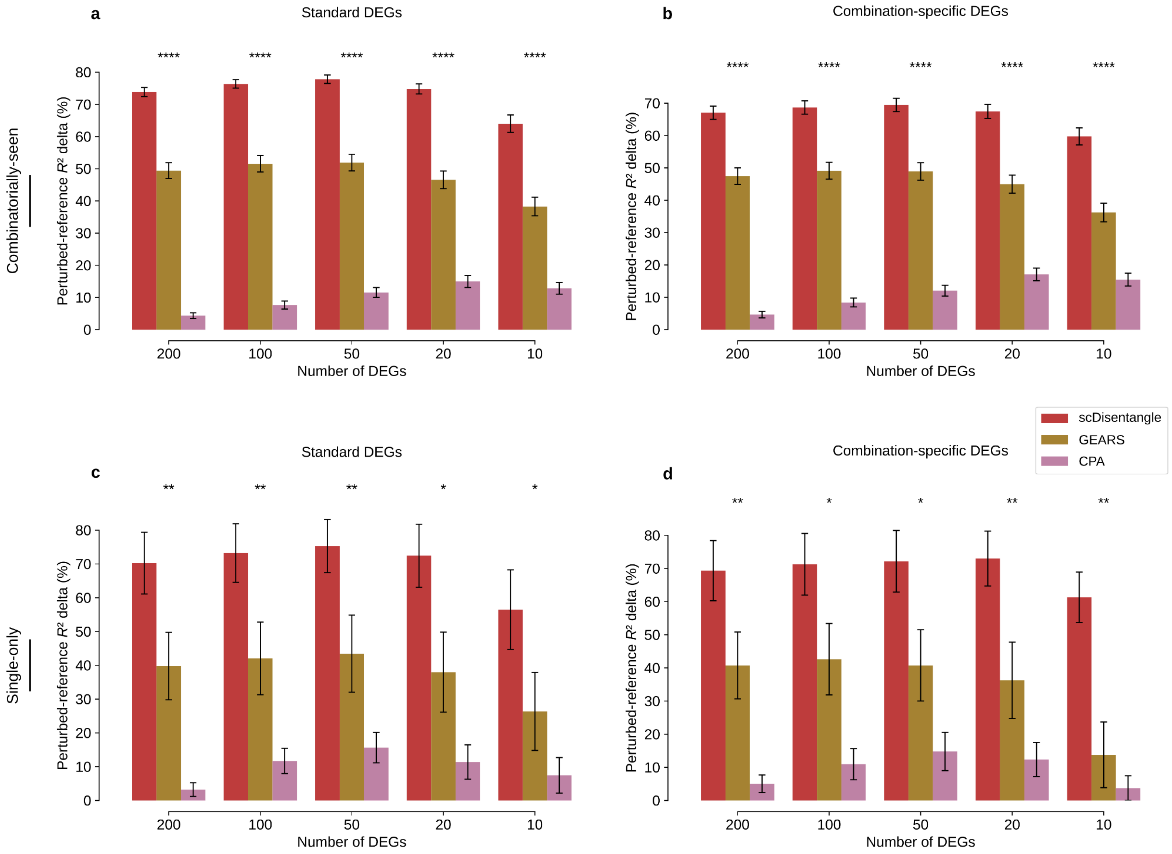


**Supplementary Fig. 8: Benchmark on Norman using Perturbed-reference $R^2$ delta.**

**a,b,** Perturbed-reference $R^2$ delta evaluated on standard DEGs (**a**) and combination-specific DEGs (**b**) across $n = 128$ evaluations (combinatorially-seen out-of-distribution scenario). **c,d,** Perturbed-reference $R^2$ delta evaluated on standard DEGs (**c**) and combination-specific DEGs (**d**) across $n = 9$ evaluations (single-only out-of-distribution scenario). Naive baselines are excluded from perturbed-reference metrics. Bars indicate the mean and error bars the standard error of the mean (s.e.m.). Significance was assessed using a one-sided paired Wilcoxon signed-rank test against the best-performing competing method at each DEG count; *$P < 0.05$, **$P < 0.01$, ***$P < 0.001$, ****$P < 0.0001$.

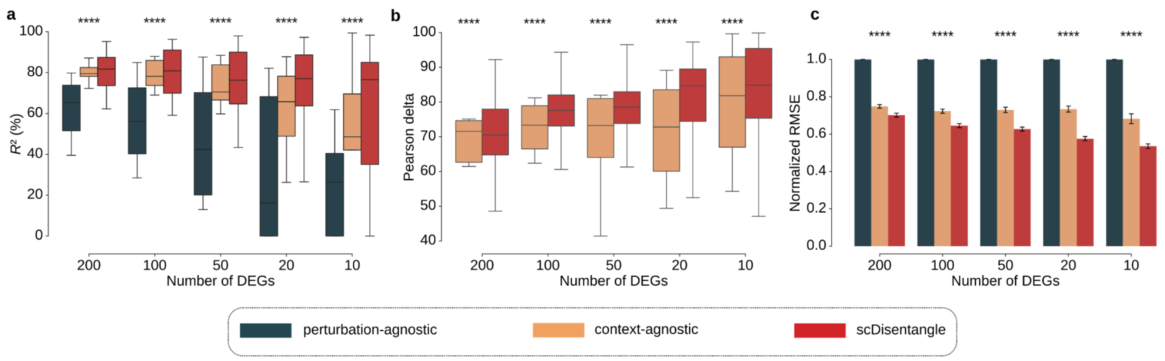


**Supplementary Fig. 9: Benchmark against naive baselines on Myocarditis (CD8+ T subset).**

**a-c,** Benchmark of scDisentangle against baselines on $R^2$ (**a**), Pearson delta (**b**), and Normalized RMSE (**c**). In boxplots (**a**,**b**), centre line indicates median, box edges the first and third quartiles, and whiskers extend to 1.5× the interquartile range; outliers not shown. In the barplot (**c**), bars indicate the mean and error bars the standard error of the mean (s.e.m.) across $n$ = 110 independent evaluations (11 OOD donors × 10 random seeds). Significance was assessed using one-sided Mann-Whitney U tests applied within each OOD context against the best-performing baseline at each DEG count; context-level p-values were combined using Fisher's method; *$P$ < 0.05, **$P$ < 0.01, ***$P$ < 0.001, ****$P$ < 0.0001.

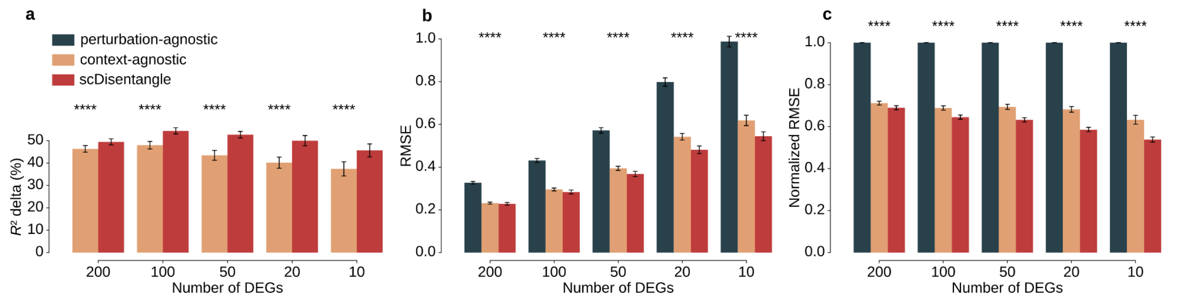


**Supplementary Fig. 10: Benchmark against naive baselines on Myocarditis (full data).**

**a-c,** Benchmark of scDisentangle against baselines on $R^2$ delta (**a**), RMSE (**b**), and Normalized RMSE (**c**). Bars indicate the mean and error bars the standard error of the mean (s.e.m.) across $n$ = 110 independent evaluations (11 OOD donors × 10 random seeds). Significance was assessed using one-sided Mann-Whitney U tests applied within each OOD context against the best-performing baseline at each DEG count; context-level p-values were combined using Fisher's method; *$P$ < 0.05, **$P$ < 0.01, ***$P$ < 0.001, ****$P$ < 0.0001.

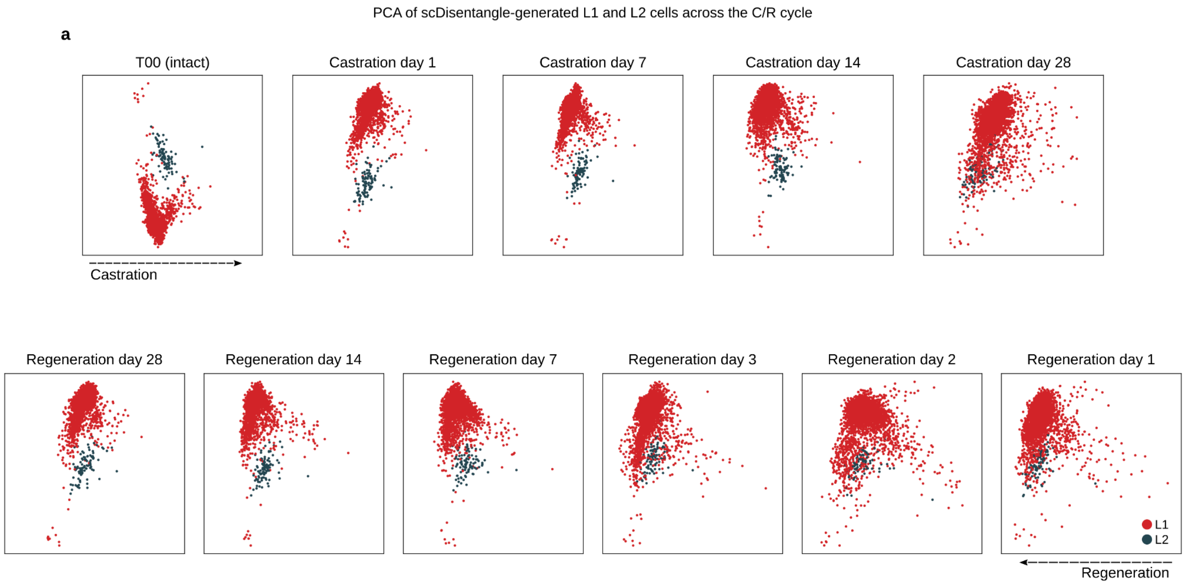


**Supplementary Fig. 11: Co-clustering and divergence of luminal L1 and L2 cells.**

**a,** PCAs of scDisentangle-generated L1 and L2 cells fitted independently at each stage coloured by cell type.

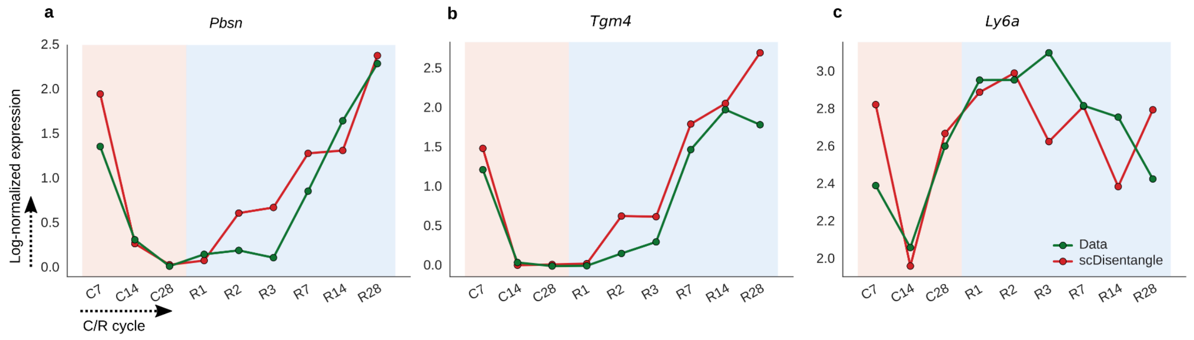


**Supplementary Fig. 12: Comparing average observed and scDisentangle-generated expression across the castration-regeneration cycle.**

**a-c,** Lineplots showing observed (green) vs. scDisentangle-generated (red) mean log-normalized expression of *Pbsn* (**a**), *Tgm4* (**b**), and *Ly6a* (**c**) in L2 cells across C/R stages (C1 excluded). These lineplots summarize violin plots of Fig. 4f.

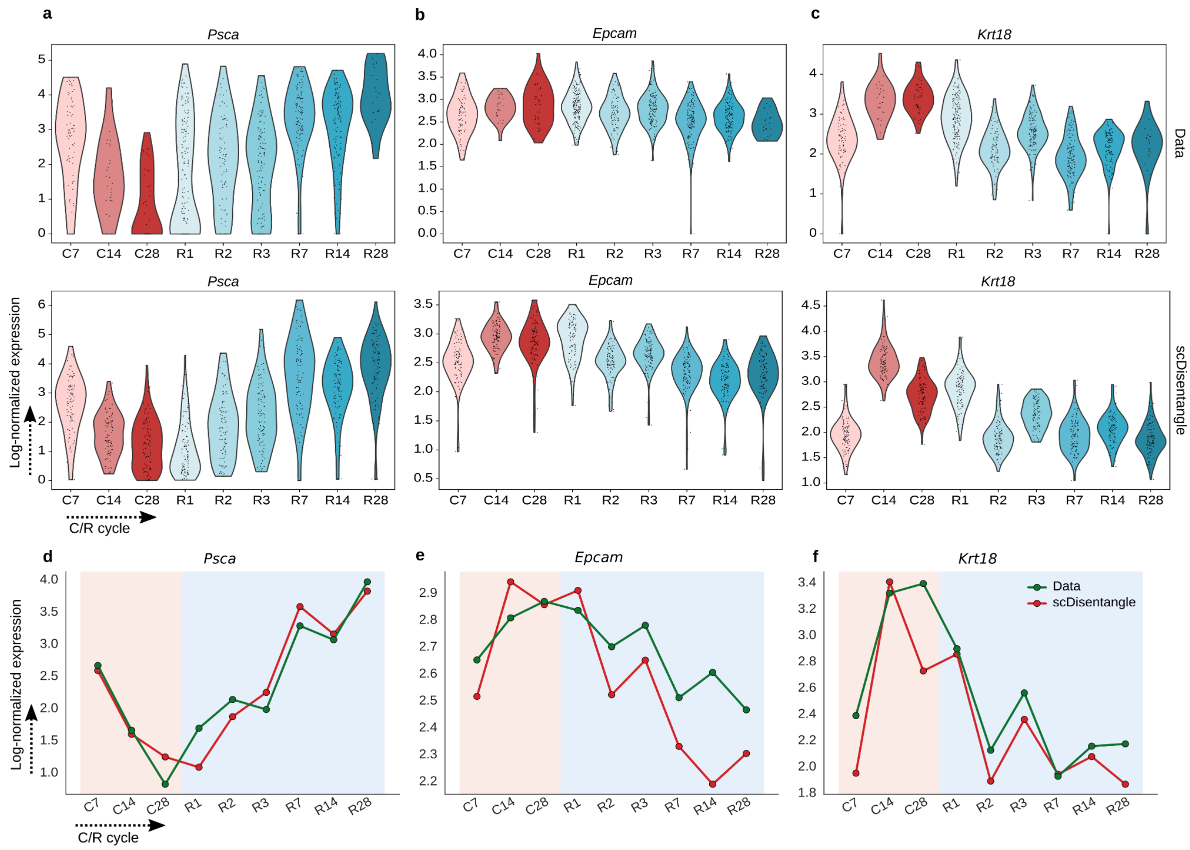


**Supplementary Fig. 13: Comparing observed and scDisentangle-generated expression for additional genes across the castration-regeneration cycle.**

**a-c,** Violin plots of observed (top row) and scDisentangle-generated (bottom row) log-normalized expression of *Psca* (**a**), *Epcam* (**b**), and *Krt18* (**c**) in L2 cells across C/R stages (C1 excluded). Individual data points overlaid. **d-f,** Lineplots showing observed (green) vs. generated (red) mean log-normalized expression of *Psca* (**d**), *Epcam* (**e**), and *Krt18* (**f**) in L2 cells across C/R stages.

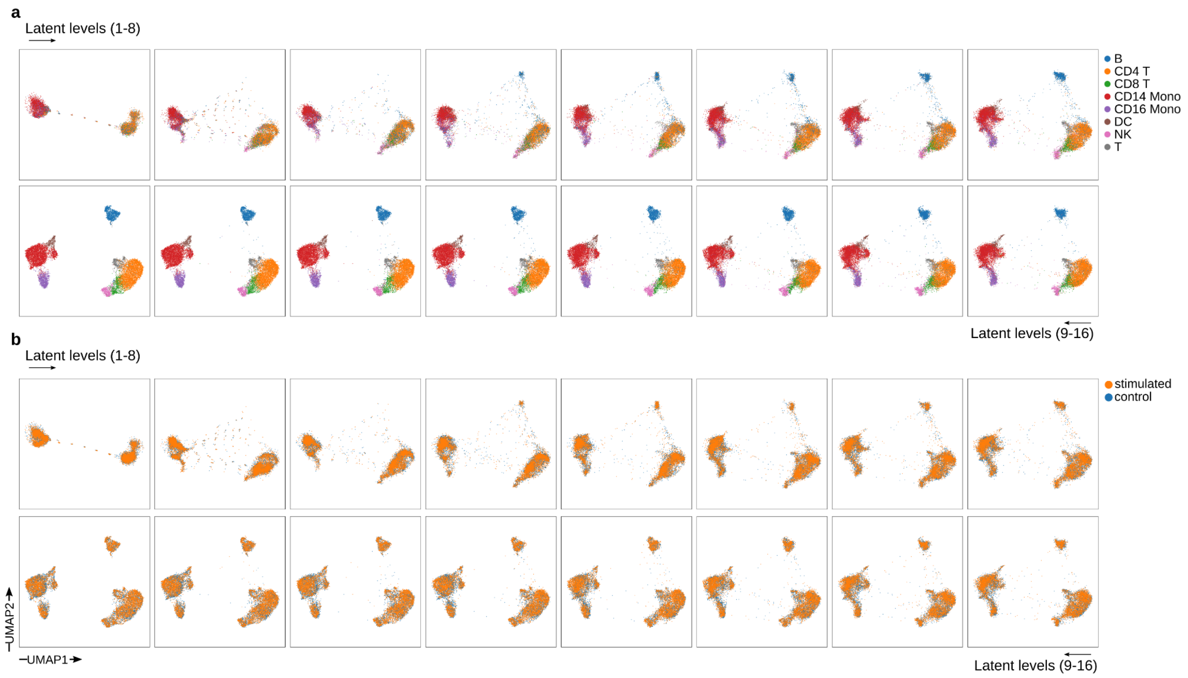


**Supplementary Fig. 14: Visualization of scDisentangle's progressive latents on Kang PBMCs. a,b,** UMAP fit on scDisentangle finest level and inferred on all levels coloured by cell type (**a**) and by condition (**b**) on Kang.

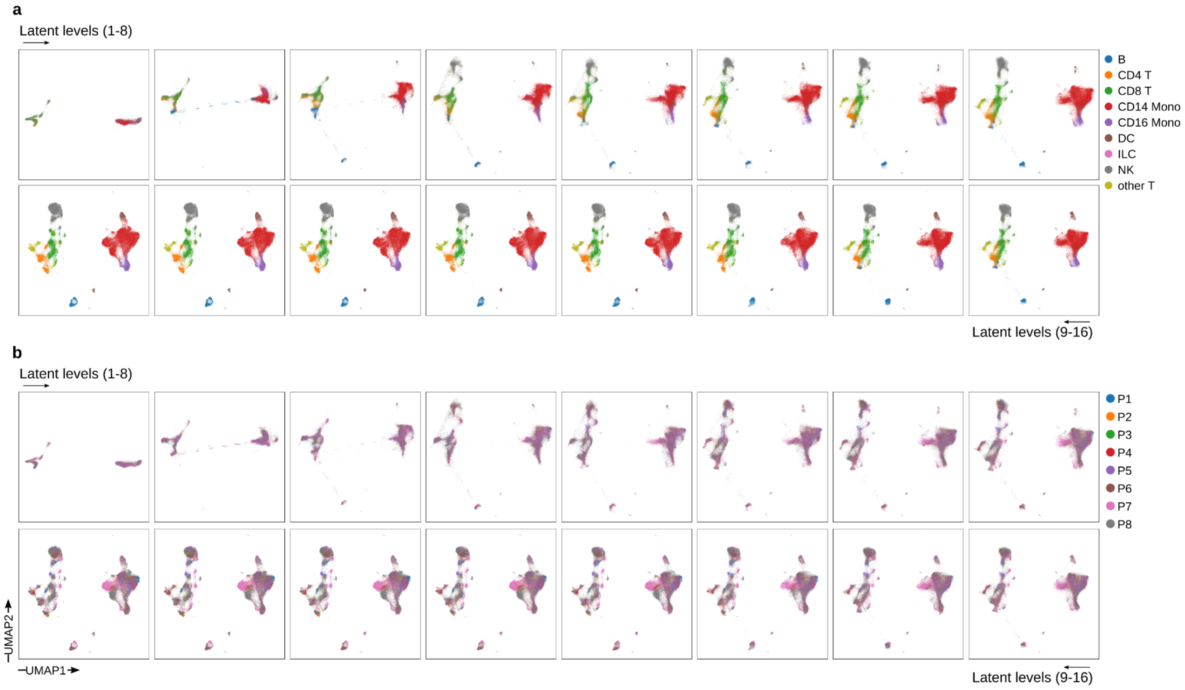


**Supplementary Fig. 15: Visualization of scDisentangle's progressive latents on Hao PBMCs.**

**a,b,** UMAP fit on scDisentangle finest level and inferred on all levels coloured by cell type (**a**) and donor (**b**) on Hao PBMC dataset.

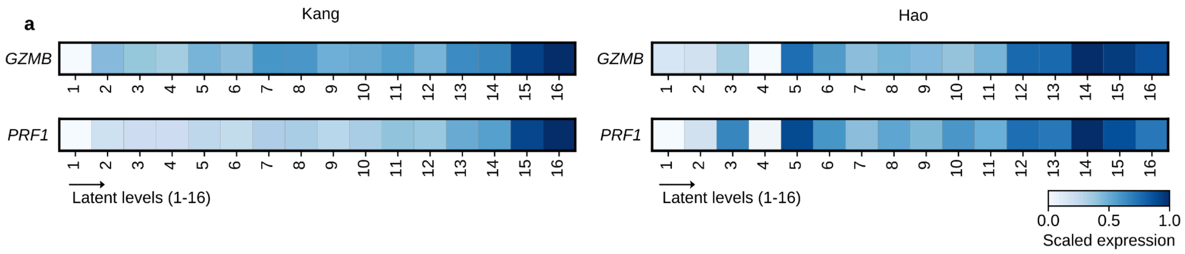


**Supplementary Fig. 16: Gene expression patterns across scDisentangle's generated levels.**

**a,** Matrixplot of mean log-normalized expression of GZMB and PRF1 per decoded latent level in CD8+ T cells on Kang (left) and Hao (right). Scaled between 0 and 1 across levels.

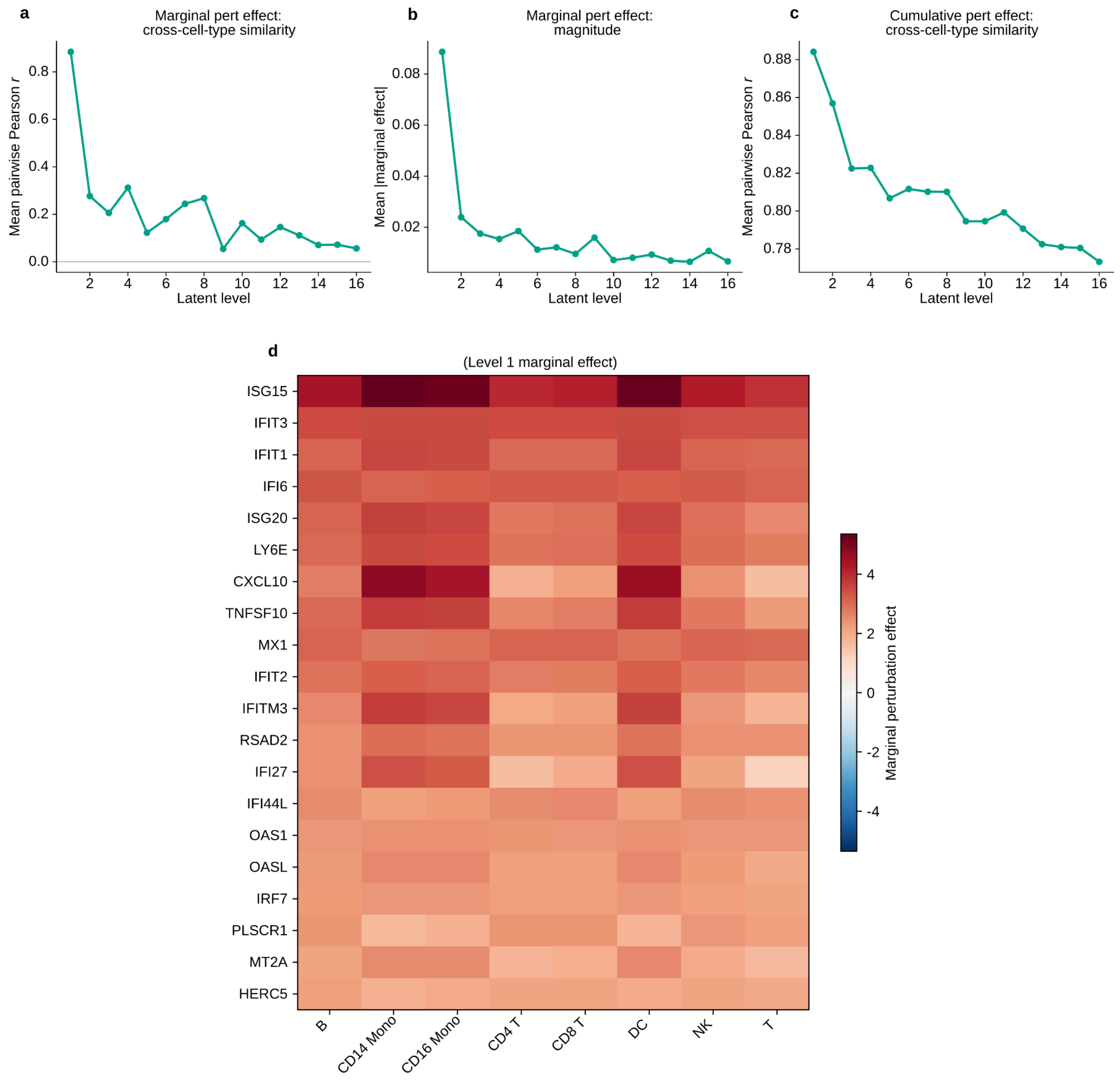


**Supplementary Fig. 17: Perturbation effect decomposition analysis on Kang.**

**a,** Mean pairwise Pearson correlation of marginal perturbation effects between all cell-type pairs ($n$ = 28 pairs from 8 cell types) at each of the 16 progressive latent levels. The marginal perturbation effect at level l is defined as the incremental contribution of that level relative to the previous level. **b,** Mean absolute marginal perturbation effect (averaged across cell types) at each level. **c,** Mean pairwise Pearson correlation of cumulative perturbation effects between cell-type pairs at each level. **d,** Heatmap of the Level 1 marginal perturbation effect for the top 20 genes ranked by absolute mean effect across all 8 cell types. Rows: genes; columns: cell types. Colour scale indicates signed marginal effect (red: higher expression upon IFN-β stimulation, blue: lower expression). All top-20 level-1 effects were positive.

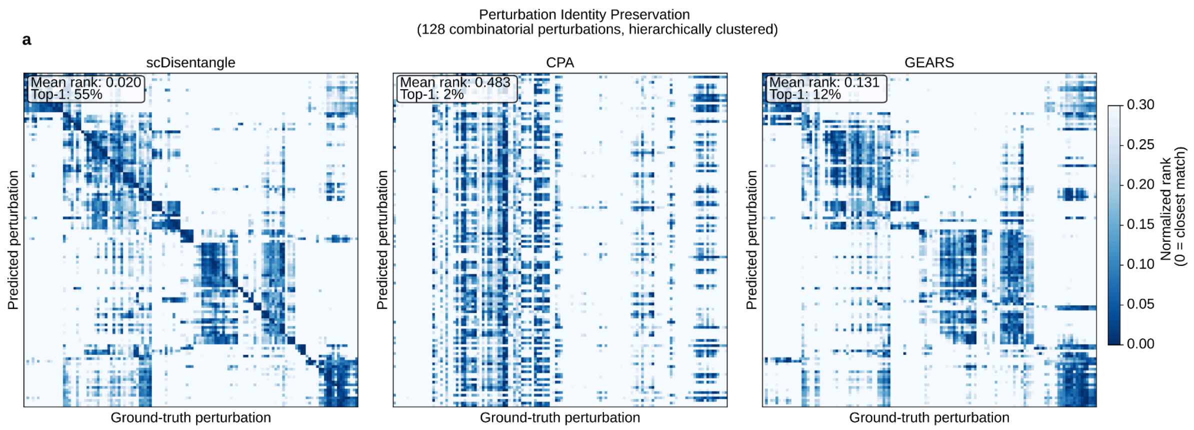


**Supplementary Fig. 18: Perturbation identity preservation for combinatorial perturbations (combinatorially-seen scenario).**

**a,** Heatmaps show the normalized rank of ground-truth perturbation centroids for each predicted perturbation, based on Euclidean distance (scDisentangle, left; CPA, middle; GEARS, right). Rows correspond to predicted perturbations; columns correspond to ground-truth perturbations. Both axes are ordered by hierarchical clustering of ground-truth centroids. Darker diagonal entries indicate that the predicted centroid is closer to its matching ground-truth centroid relative to other ground-truth centroids. Mean rank denotes the average normalized rank of the matching ground-truth perturbation across all predicted perturbations; Top-1 denotes the percentage of predicted perturbations for which the matching ground-truth centroid is the closest ground-truth centroid.

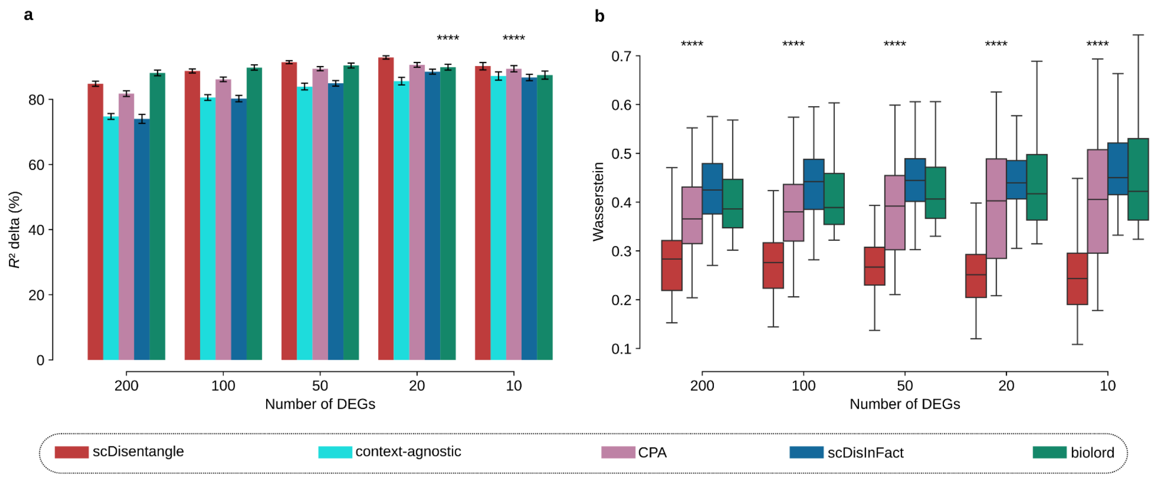


**Supplementary Fig. 19: Benchmark on Liver spatio-temporal experiment.**

**a,** $R^2$ delta. **b,** Wasserstein distance. Models are trained 10 seeds per OOD setting, and evaluated on timepoint-specific infection response DEGs (n = 100 evaluations). scGen excluded. Bars (**a**) indicate the mean and error bars the standard error of the mean (s.e.m.). In the boxplot (**b**) centre line indicates median, box edges the first and third quartiles, and whiskers extend to 1.5× the interquartile range; outliers not shown. Significance was assessed using one-sided Mann-Whitney U tests applied within each OOD context against the best-performing competing method at each DEG count; context-level p-values were combined using Fisher's method; *$P$ < 0.05, **$P$ < 0.01, ***$P$ < 0.001, ****$P$ < 0.0001. Stars are shown only when the median difference also favours scDisentangle.
